# The CMG helicase bypasses DNA protein cross-links to facilitate their repair

**DOI:** 10.1101/381582

**Authors:** Justin L. Sparks, Alan O. Gao, Markus Räschle, Nicolai B. Larsen, Matthias Mann, Julien P. Duxin, Johannes C. Walter

## Abstract

Covalent and non-covalent nucleoprotein complexes impede replication fork progression and thereby threaten genome integrity. Using *Xenopus laevis* egg extracts, we previously showed that when a replication fork encounters a covalent DNA-protein cross-link (DPC) on the leading strand template, the DPC is degraded to a short peptide, allowing its bypass by translesion synthesis polymerases. Strikingly, we show here that when DPC proteolysis is blocked, the replicative DNA helicase (CMG), which travels on the leading strand template, still bypasses the intact DPC. The DNA helicase RTEL1 facilitates bypass, apparently by translocating along the lagging strand template and generating single-stranded DNA downstream of the DPC. Remarkably, RTEL1 is required for efficient DPC proteolysis, suggesting that CMG bypass of a DPC normally precedes its proteolysis. RTEL1 also promotes fork progression past non-covalent protein-DNA complexes. Our data suggest a unified model for the replisome’s response to nucleoprotein barriers.

## Introduction

DNA replication forks encounter numerous obstacles that challenge the process of genome duplication. Discrete DNA lesions (e.g. pyrimidine dimers, oxidized bases, abasic sites) generally stall the replicative DNA polymerase but not the helicase, leading to helicase-polymerase uncoupling (Byun et al., 2005). Bulkier obstacles also block the replicative DNA helicase and therefore stall the entire replisome. These include DNA interstrand cross-links (ICLs; (Fu et al., 2011)) and DNA protein cross-links (DPCs; (Duxin et al., 2014)), in which a protein is covalently linked to one strand of the DNA. DPCs are formed by exogenous agents such ultraviolet light and ionizing radiation, as well as the chemotherapeutics cisplatin, mitomycin C, and nitrogen mustard (Ide et al., 2011; Stingele et al., 2017; Vaz et al., 2017). DPCs are also generated by endogenous agents such as formaldehyde (a byproduct of histone and DNA demethylation), abasic sites (generated from DNA glycosylases or spontaneous hydrolysis), reactive oxygen species, and topoisomerases. In addition to ICLs and DPCs, DNA binding proteins that are tightly bound to DNA interfere with replication fork progression. These non-covalent nucleoprotein complexes include RNA polymerases, tightly bound transcription factors, centromeric chromatin, and pre-replication complexes. Here, we address how the vertebrate replisome overcomes covalent and non-covalent nucleoprotein complexes.

Recent studies have greatly advanced our understanding of how eukaryotic organisms repair covalent DPCs. DPCs smaller than ~8 kD can be excised by nucleotide excision repair whereas larger DPCs require more complex pathways (Ide et al., 2011). While some studies proposed that the response to large DPCs involves strand breakage and recombination, this idea has been challenged (Rosado et al., 2011). In 2014, Wss1 was identified as a DNA-dependent protease that degrades DPCs in yeast, and genetic experiments suggested it operates in S phase (Stingele et al., 2014). Contemporaneous experiments in *Xenopus laevis* egg extracts showed that when a replication fork collides with a DPC, the DPC undergoes proteolysis by an unknown protease, leaving behind a short peptide adduct that is bypassed by translesion DNA polymerases (Figure 1A) (Duxin et al., 2014). Together, these studies established the existence of a protease-mediated DPC repair pathway that avoids the need for strand cleavage and recombination.

**Figure 1.**
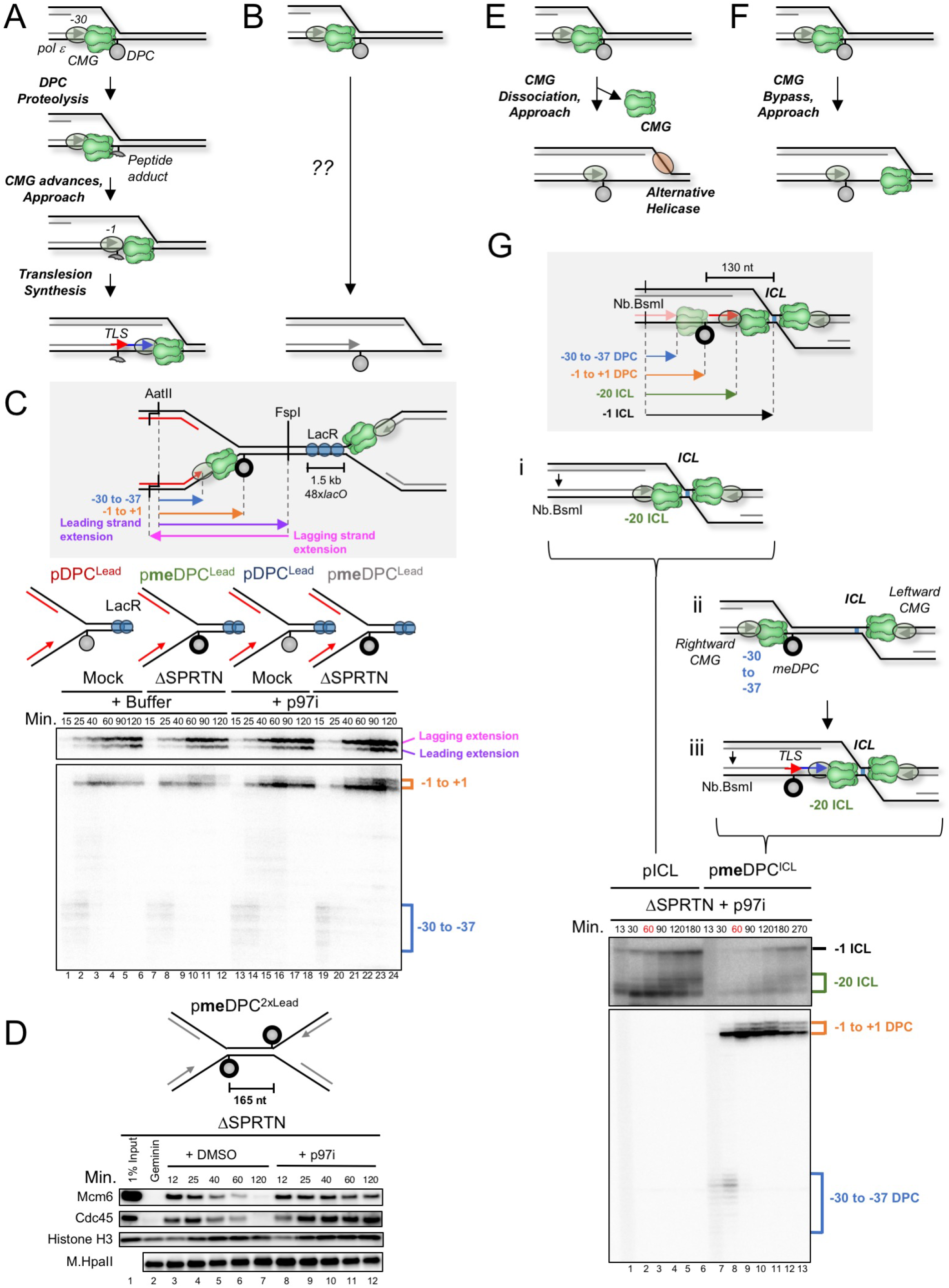
CMG bypasses an intact DNA-protein crosslink on its translocation strand. (**A**) Current model of replication-coupled DPC repair (Duxin *et al*. 2014). (**B**) Schematic of what happens in the presence of Ub-VS (Duxin *et al*. 2014). (**C**) Grey inset: Schematic of nascent leading strand products released by AatII and FspI digestion of pmeDPC^Lead^ or pDPC^Lead^. pDPC^Lead^ or pmeDPC^Lead^ were prebound with LacR to prevent one replication fork from colliding with the DPC. The plasmids were replicated in mock-depleted or SPRTN-depleted egg extract containing ^32^P[α]-dATP and supplemented with DMSO or p97 inhibitor (p97i). At different times, DNA was recovered and digested with AatII and FspI, separated on a denaturing polyacrylamide gel, and visualized by autoradiography. The lower autoradiogram shows nascent leading and lagging strands generated by the rightward replication fork, and the uppper autoradiogram shows extension products. The CMG footprint is bracketed in blue (−30 to −37), and the products stalled at the adducted base are bracketed in orange (−1 to +1). (**D**) pmeDPC^2xLead^ was replicated in SPRTN-depleted egg extract supplemented with DMSO or p97i (NMS-873). At different times, plasmid-associated proteins were recovered and blotted with the indicated antibodies. Samples were also withdrawn and processed by the stringent pull-down procedure described in Figure S1D for the M.HpaII panel. (**E-F**) Alternative models to explain the disappearance of the CMG footprint at stable DPCs: CMG dissociation (E) or CMG bypass (F). (**G**) Grey inset: nascent leading strand products released by nicking with Nb.BsmI on pmeDPC^ICL^ before (transparent CMG) and after (solid CMG) collision of CMG with the ICL. pICL or pmeDPC^ICL^ was replicated in SPRTN-depleted egg extract supplemented with p97i to prevent CMG unloading at the ICL. After 60 minutes (red), when leading strands have approached the DPC, an equal volume of fresh HSS supplemented with p97i was added to enhance translesion synthesis past the DPC. At different times, DNA was recovered and nicked with Nb.BsmI (to specifically release only the nascent leading strands), separated on a denaturing polyacrylamide gel, and visualized with autoradiography. The CMG footprint at the DPC is bracketed in blue (−30 to −37), the approach product at the DPC in orange (−1 to +1), the CMG footprint at the ICL in green (−20), and the approach at the ICL in black (−1).

Like Wss1, its vertebrate ortholog SPRTN (also named Spartan or DVC1) is a DNA-dependent protease (Lopez-Mosqueda et al., 2016; Morocz et al., 2017; Stingele et al., 2016; Vaz et al., 2016). In mammalian cells expressing protease-deficient SPRTN, DPCs accumulate on chromatin, replication fork progression is severely impaired, and DNA damage and genome instability are observed (Lessel et al., 2014; Maskey et al., 2017; Maskey et al., 2014; Vaz et al., 2016). In humans, mutations in SPRTN’s protease domain cause Ruijs-Aalfs syndrome (RJALS), a human genetic disease characterized by genome instability, premature aging, and early onset liver cancer (Lessel et al., 2014). Collectively, these experiments suggest that SPRTN suppresses cell death, liver cancer, and aging by removing DPCs that block replication fork progression. In frog egg extracts, SPRTN and the proteasome have overlapping functions in promoting replication-coupled DPC proteolysis (Gao et al., submitted; manuscript enclosed). In this system, SPRTN activity requires that a nascent strand be extended to within a few nucleotides of the DPC, whereas proteasome activity requires DPC ubiquitylation and the presence of ssDNA near the adduct. Despite this progress, important questions remain. For example, given that proteolysis is triggered by the collision of a DNA replication fork with the DPC (Duxin et al., 2014), it is unclear how inadvertent destruction of the replisome itself is avoided during DPC repair.

The replicative DNA helicase is the first component of the replisome to encounter nucleoprotein complexes on the DNA template and understanding how the helicase overcomes such obstacles is critical. All replicative helicases that have been studied in detail form hexameric rings that motor along one of the two template strands using the energy harnessed from ATP hydrolysis, leading to DNA unwinding (“steric exclusion model”)(O’Donnell and Li, 2018). The bacterial replicative DnaB helicase, which translocates with 5’ to 3’ polarity along the lagging strand template, stalls when it encounters non-covalent nucleoprotein complexes. To motor past these obstacles, it employs two accessory helicases, UvrD and Rep, which act redundantly (Guy et al., 2009). Rep interacts directly with DnaB and is therefore more potent than UvrD in overcoming obstacles. Importantly, Rep and UvrD are 3’ to 5’ helicases and thus have the opposite polarity to DnaB. As such, they assemble on the leading strand template and likely cooperate with DnaB in overcoming obstacles.

In contrast to DnaB, the eukaryotic replicative DNA helicase, CMG (a complex of Cdc45, MCM2-7, and GINS), encircles and translocates 3’ to 5’ along the leading strand template (Fu et al., 2011). While purified CMG stalls at a biotin-streptavidin (SA) complex on the lagging strand template, MCM10 helps overcome the barrier (Langston and O’Donnell, 2017; Langston et al., 2017). It was proposed that binding by MCM10, which lacks known enzymatic activity, alters CMG’s interaction with the lagging strand to allow bypass. In contrast, MCM10 does not promote efficient CMG bypass of a leading strand biotin-SA complex, suggesting that CMG cannot translocate past a bulky leading strand adduct ((Langston et al., 2017) see also (Nakano et al., 2013)). Interestingly, the large T antigen DNA helicase can by itself bypass a DPC on the translocation strand, perhaps via transient ring-opening (Yardimci et al., 2012). Whether CMG progression past covalent or non-covalent nucleoprotein complexes is assisted by accessory helicases, as seen in bacteria, has not been directly examined. However, consistent with this idea, the 5’ to 3’ helicases Rrm3 and Pif1 help replication fork progression past non-covalent nucleoprotein complexes in yeast (Ivessa et al., 2000). These results suggest that, like bacteria, yeast cells overcome obstacles by engaging an accessory DNA helicase that moves on the strand opposite the one hosting the replicative DNA helicase.

Vertebrates contain at least six 5’ to 3’ DNA helicases (XPD, DDX3, DDX11/CHLR1, FANCJ/BRIP/BACH1, PIF1, and RTEL1) that could cooperate with CMG in the response to replication obstacles (Brosh, 2013). XPD participates in nucleotide-excision repair and has no known role in fork dynamics. DDX3 unwinds DNA, but nothing is known about its role in genome maintenance (Garbelli et al., 2011). DDX11, FANCJ, and PIF1 have been implicated in genome maintenance through roles in cohesion establishment, ICL repair, and replication fork progression (Cali et al., 2016; Cantor and Nayak, 2016; Gagou et al., 2014). Deletion of RTEL1 in mice is lethal, and hypomorphic mutations in the human gene cause Hoyeraal-Hreidarsson syndrome, which is characterized by telomere shortening, bone marrow failure, immunodeficiency, growth retardation, and microcephaly (Vannier et al., 2014). While RTEL1 was first identified as a regulator of telomere homeostasis, it also promotes replication fork progression through an unknown mechanism (Vannier et al., 2013). In summary, vertebrate cells contain several potential accessory replicative helicases, but none have been specifically implicated in overcoming nucleoprotein complexes.

Based on experiments in *Xenopus* egg extracts, we previously proposed that DPC proteolysis is essential for the CMG helicase to move past a DPC (Figure 1A)(Duxin et al., 2014). We now show that CMG can bypass a DPC on the translocation strand in the absence of DPC proteolysis. Efficient bypass requires RTEL1, which probably translocates along the lagging strand template to unwind DNA past the DPC. Importantly, RTEL1 is also required for efficient DPC proteolysis. Therefore, our data suggest that when a replication fork encounters a DPC on the leading strand template, CMG first bypasses the adduct with the assistance of RTEL1, followed by DPC proteolysis and translesion synthesis (TLS) across the lesion. RTEL1 also promotes fork progression past non-covalent nucleoprotein complexes. Our data support a unified model that explains how accessory helicases promote replicative bypass of nucleoprotein complexes in eukaryotic organisms.

## Results

### Leading strands rapidly approach a non-degradable DPC

To study the mechanism of DPC repair, we incubated the 45 kDa DNA methyltransferase M.HpaII with a plasmid containing a fluorinated M.HpaII recognition site (CCGG), which leads to covalent trapping of the enzyme on DNA (Figure S1A; (Chen et al., 1991; Duxin et al., 2014)). We previously showed that when this plasmid (pDPC) is replicated in *Xenopus* egg extracts, leading strands stall 30 nucleotides (nt) from the adducted base due to the footprint of the CMG helicase, which translocates along the leading strand template (Figure 1A; (Duxin et al., 2014)). After a short pause, synthesis resumes and nascent leading strands approach to within 1 nucleotide of the DPC (“approach”), which they subsequently bypass using the REV1-DNA pol ζ complex (Figure 1A, (Duxin et al., 2014)). Because degradation of the DPC roughly coincides with the disappearance of the 30 nt CMG footprint, the results suggested that degradation of the DPC allows CMG to move beyond the adduct (Duxin et al., 2014)(Figure 1A). Surprisingly, however, when DPC proteolysis is blocked via ubiquitin depletion (by supplementing extracts with the de-ubiquitylating enzyme inhibitor Ub-VS), the nascent leading strand still approaches the adduct, albeit very inefficiently, suggesting that CMG is cleared from its position in front of the DPC adduct (Figure 1B; (Duxin et al., 2014)). Moreover, the nascent lagging strand is ultimately extended beyond the DPC. These results show that the replication fork can eventually move past an intact DPC, but the fate of CMG in this process was unclear (Figure 1B).

Given the importance of ubiquitin signaling in the DNA damage response, we wanted to block degradation of the DPC without ubiquitin depletion and examine the effect on approach. To this end, we took advantage of the fact that the DPC is degraded in egg extracts by two overlapping, replication-coupled pathways (Gao et al., submitted, manuscript enclosed). In one pathway, arrival of the replication fork at the DPC stimulates DPC proteolysis by SPRTN (Figure S1B). In the other pathway, replication-dependent ubiquitylation of the DPC promotes its destruction by the proteasome (Figure S1B). To inhibit the SPRTN pathway, we used egg extract immunodepleted of SPRTN (Figure S1C); to inhibit the proteasome pathway, we employed a DPC whose lysine residues had been chemically methylated to prevent ubiquitylation (meDPC; Figure 1C, spheres with bold outlines). Under these conditions, the DPC was not degraded, and we refer to this as a “stable” DPC (Figure S1D, lanes 7-11, upper panel; (Gao et al., submitted)). To ensure that a single fork encountered the stable DPC on the leading strand template, we flanked the meDPC on the right with a 48x*lacO* array bound by Lac repressor (LacR), which blocks arrival of the leftward, converging fork (see grey inset in Figure 1C; (Duxin et al., 2014)). To monitor progress of the rightward leading strand, we digested the DNA with AatII and FspI (Figure 1C; top grey inset). Strikingly, in the absence of ubiquitin depletion, the 30 nt CMG footprint disappeared with the same kinetics as when proteolysis was unimpeded (Figure 1C; compare lanes 1-6 and 7-12; Figure S1E for quantification). The staggered cuts made by AatII also allowed us to distinguish leading and lagging strand extension products (Figure 1C; grey inset, pink and purple arrows), which revealed that the lagging strand was readily extended past the stable DPC (Figure 1C, lanes 7-12, top autoradiogram). Compared to the lagging strand, leading strand extension past the stable DPC was severely reduced, though not eliminated, consistent with defective TLS past the adduct. Collectively, these results show that when ubiquitin levels are normal, the CMG footprint disappears from the DPC with the same kinetics whether or not the DPC undergoes proteolysis, and the lagging strand is extended past the DPC. There are two possible interpretations of this result. First, upon fork stalling, CMG rapidly dissociates from chromatin, thereby allowing leading strand approach to the lesion even without DPC proteolysis (Figure 1E). In this scenario, a helicase other than CMG unwinds DNA past the DPC to allow lagging strand extension (Figure 1E, orange sphere). Second, CMG bypasses the stable DPC (Figure 1F). If CMG normally bypasses the DPC before its destruction, leading strand approach and lagging strand extension would also be unaffected by DPC proteolysis.

### CMG bypasses a stable DPC

To address whether CMG dissociates from or translocates past a stable DPC, we examined approach in the presence of an inhibitor of the p97 ATPase (p97i), which is required to extract CMG from chromatin during replication termination and ICL repair (Dewar et al., 2017; Fullbright et al., 2016; Maric et al., 2014; Moreno et al., 2014; Semlow et al., 2016). If CMG has to dissociate upon stalling at the DPC to enable approach (Figure 1E), p97i should delay approach. In contrast, if CMG bypasses the DPC to enable approach (and dissociates only later; Figure 1F), the p97i should have no effect. As shown in Figure 1C, p97i did not slow the kinetics of leading strand approach to or lagging strand extension past a stable DPC (compare lanes 7-12 and 19-24; Figure S1E for quantification of approach). Similarly, p97i had no effect on approach when DPC proteolysis was inhibited with Ub-VS (Figure S1F). To directly examine the fate of CMG after DPC encounter and how this is affected by p97i, we monitored CMG binding to the plasmid pmeDPC^2xLead^ in SPRTN-depleted extract, where both converging CMGs encounter a stable DPC on the leading strand template (Figure 1D; cartoon). As shown in Figure 1D, CMG gradually dissociated from this plasmid, but not when p97i was included in the reaction (see Figure S1G for the analogous result with Ub-VS). The data show that p97i completely blocked unloading of CMGs from a DPC-containing plasmid. However, given that the kinetics with which the CMG footprint disappeared at a stable DPC was completely unaffected by p97i (Figure 1C; Figure S1E for quantification), CMG must have bypassed the DPC (Figure 1F). We infer that the CMG unloading observed in the absence of p97i results from CMG bypass, convergence with another CMG, and removal via the mechanism employed during replication termination (Figure S1H; (Dewar et al., 2015; Dewar et al., 2017)).

To further test whether CMG bypasses a stable DPC, we sought to trap CMG on the other side of a DPC after bypass. To this end, we created a new construct, pmeDPC^ICL^, in which a meDPC was flanked on the right by an ICL (Figure 1G). When replicated in ΔSPRTN extract containing p97i to prevent CMG unloading at the ICL, the leftward moving CMG can readily reach the ICL, whereas the rightward CMG must bypass the stable DPC before it can reach the ICL (Figure 1Gii-iii). After CMG bypass and TLS, the characteristic −20 footprint that is observed when a CMG collides with an ICL (Raschle et al., 2008) should be generated by the rightward leading strand (Figure 1Giii). As shown in Figure 1G, lower autoradiogram, the CMG footprint at the DPC disappeared by 60 minutes (lane 9; evident from the loss of −30 to −37 products), when the DPC was still completely intact (Figure S1I). With time, the −20 ICL footprint of the rightward leading strand gradually accumulated at the ICL, indicating that the rightward CMG had bypassed the DPC and reached the ICL (Figure 1G upper autoradiogram; “−20”). The slow appearance of the footprint and its relatively low abundance reflect the low efficiency of leading strand TLS past an intact DPC (Figure 1C, lanes 7-12, upper autoradiogram). We were also able to detect the CMG footprint downstream of a DPC that was stabilized with Ub-VS (Figure S1J). DPC bypass was not affected when an inhibitor of Dbf4-dependent kinase was added immediately after forks had reached the DPC, indicating that bypass does not involve new origin firing downstream of the DPC (Figure S1K). Together, our data strongly support the idea that CMG is able to bypass a bulky DPC on the translocation strand. Given that there is no known pathway to load MCM2-7 *de novo* in S phase, and because the CMG footprint disappears in the absence of CMG dissociation, we infer that the same molecule of MCM2-7 that initially collides with the DPC as part of CMG undergoes bypass.

### ssDNA downstream of a stable DPC facilitates CMG bypass

When the bacterial replicative helicase DnaB encounters nucleoprotein complexes, accessory helicases that translocate on the strand opposite the one encircled by DnaB help the replisome bypass the obstacle (Guy et al., 2009). Inspired by this model, we speculated that a 5’ to 3’ accessory helicase might load onto the lagging strand template and assist CMG bypass of a leading strand DPC (Figure 2Ai). As an initial test of this idea, we placed a second meDPC on the lagging strand template 15 nt upstream of the leading strand DPC (pmeDPC^Lag/Lead^), as this might block unwinding by a 5’ to 3’ helicase (Figure 2Aii). We also placed the second DPC 15 nt downstream of the leading strand DPC (pmeDPC^Lead/Lag^), reasoning that this would allow a 5’ to 3’ helicase to unwind some DNA at the DPC (Figure 2Aiii). Consistent with the action of a 5’ to 3’ helicase, CMG bypass (as measured via disappearance of the CMG footprint) was severely inhibited by the upstream but not the downstream lagging strand DPC (Figure 2B, compare lanes 19-24 with 25-30; see graph for quantification). This model further predicts that a ssDNA bubble placed immediately downstream of the DPC should rescue CMG bypass on pmeDPC^Lag/Lead^ (Figure 2Aiv; pmeDPC^Lag/Lead-Bubble^). Since there are only 4 bp between the DPC attachment site and the bubble, arrival of CMG should be sufficient to melt the duplex surrounding the DPC (Figure 2Aiv). As shown in Figure 2B (lanes 31-36), a 40 nt ssDNA bubble fully rescued bypass in the presence of the upstream lagging strand DPC. Remarkably, bypass was now even faster than on the DNA template lacking any lagging strand obstruction (Figure 2B, compare lanes 31-36 with lanes 7-12). Similar results were observed when a DPC was stabilized with Ub-VS (Figure S2). Interestingly, the presence of two tandem meDPCs on the leading strand template also severely impaired bypass (Figure 2B, lanes 13-18), suggesting that a single CMG cannot simultaneously accommodate two DPCs during bypass. Together, the data suggest that generation of ssDNA downstream of the lesion is rate-limiting for CMG bypass of an intact leading strand DPC, consistent with assistance by a 5’ to 3’ helicase.

**Figure 2.**
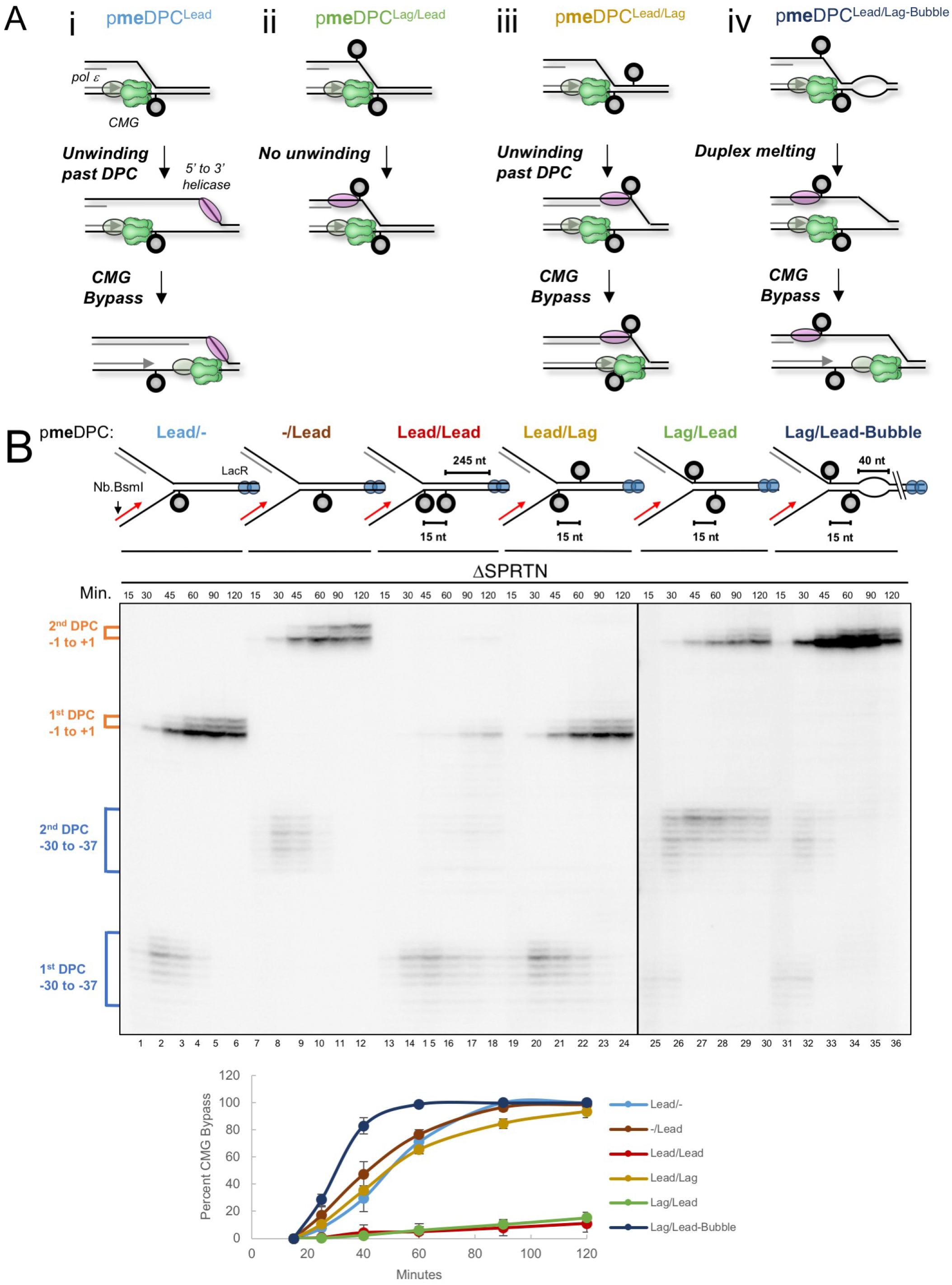
ssDNA downstream of an intact DPC facilitates CMG bypass. (**A**) Proposed action of a 5’ to 3’ helicase (pink) on four substrates (i-iv). (**B**) pmeDPC^Lead/−^, pmeDPC^−/Lead^, pmeDPC^Lead/Lead^, pmeDPC^Lead/Lag^, pmeDPC^Lag/Lead^, or pmeDPC^Lag/Lead-Bubble^ were prebound with LacR, replicated in SPRTN-depleted egg extracts, and analyzed as in Figure 1G. Disappearance of the CMG footprint (approach) was used as a proxy for CMG bypass and quantified as in Figure 1C (see methods). The mean of three experiments is shown. Error bars represent the standard deviation.

### RTEL1 promotes efficient CMG bypass of a stable DPC

Vertebrates contain at least six 5’ to 3’ DNA helicases (RTEL1, FANCJ, PIF1, DDX3, DDX11, and XPD). Using plasmid pull-down combined with mass spectrometry (PP-MS), all of these helicases except DDX11 were detected on chromatin in egg extracts (Figure 3A; (Gao et al., submitted)). Two of the helicases (PIF1 and RTEL1) bound to pDPC and an undamaged plasmid (pCTRL) in a replication-dependent fashion, suggesting that these helicases travel with the replication fork. However, the levels of RTEL1 increased between the 12 and 20-minute time points, suggesting that after fork stalling, there may be additional RTEL1 recruitment (see also Figure 5E). FANCJ and DDX3 were specifically enriched on DPC-containing plasmid, whereas XPD bound independently of replication. Depletion of PIF1, FANCJ, or DDX3 had no significant effects on bypass of a stable DPC (data not shown; we did not examine XPD or DDX11). In contrast, immunodepletion of RTEL1 (Figure 3B; see Figure S3A for explanation of RTEL1 isoforms in lane 1 of Figure 3B) greatly delayed approach of leading strands to a stable DPC (Figure 3C), and it delayed the advance of the lagging strand beyond the DPC (Figure 3D, lanes 6-10, upper autoradiogram). These results indicate that RTEL1 is required for CMG bypass and that bypass allows new Okazaki fragment priming downstream of the adduct (Figure S3B). RTEL1 depletion had no effect on the overall efficiency of DNA synthesis (Figure S3C). The defect in CMG bypass was partially rescued by wild type recombinant RTEL1 but not by an ATPase deficient mutant of RTEL1 (RTEL1^K48R^), which instead further inhibited bypass (Figure 3C; Figure S3D). Importantly, RTEL1 depletion had only a modest effect on bypass when a single-stranded DNA bubble was placed downstream of the DPC, and it did not further inhibit bypass when a lagging strand meDPC was present upstream (Figure S3E). Collectively, these data indicate that RTEL1 enables CMG to efficiently bypass intact DPCs on the leading strand template, primarily by generating a short patch of ssDNA near the adduct. The residual bypass observed in RTEL1-depleted extracts (Figure 3C) could be due to incomplete RTEL1 depletion, the presence of partially compensating accessory helicases, or bypass that is independent of accessory helicases.

**Figure 3.**
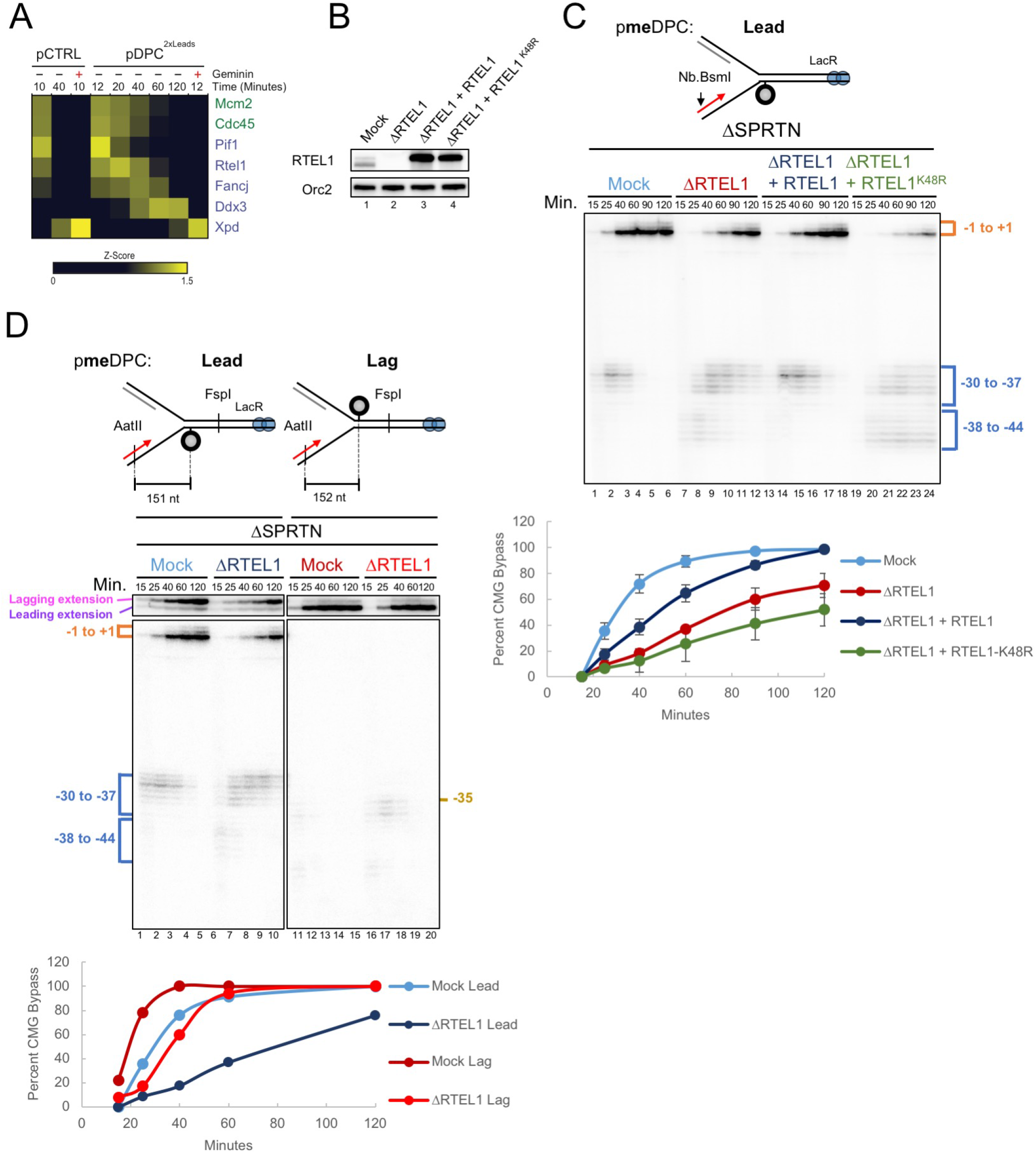
RTEL1 is required for efficient CMG bypass. (**A**) Recovery of 5’ to 3’ helicases in the mass spectrometry dataset of Gao et al. (submitted). The heat map shows the relative abundance of the indicated proteins (expressed as a z-score with yellow indicating higher abundance) for each of the conditions indicated. To identify proteins whose binding is replication-dependent, pCTRL and pDPC were incubated in extracts containing geminin, a replication licensing inhibitor. (**B**) Mock-depleted, RTEL1-depleted, and RTEL1-depleted egg extracts supplemented with wild-type RTEL1 or RTEL1-K48R were blotted with RTEL1 and ORC2 (loading control) antibodies. (**C**) Egg extracts were depleted of SPRTN and RTEL1, as indicated, and supplemented with buffer, wild-type RTEL1, or RTEL1-K48R. Leading strand approach was visualized and quantified as in Figure 1G. The mean of three experiments is shown. Error bars represent the standard deviation. (**D**) pmeDPC^Lead^ or pmeDPC^Lag^ was replicated in the indicated extracts and analyzed as in Figure 1C. As for the −38 arrest seen with leading strand M.HpaII, we ascribe the −35 arrest to collision of CMG with lagging strand M.HpaII whose underlying DNA has not been unwound. The three nucleotide difference probably reflects asymmetry in the collision of CMG with leading and lagging strand M.HpaII. CMG bypass was quantified as in Figure 1C. Figure S3F shows a repetition of this experiment without quantification.

The structures of DNA methyltransferases bound to DNA (Klimasauskas et al., 1994; Reinisch et al., 1995) suggest that M.HpaII likely interacts intimately with both strands of the double helix. Therefore, covalent coupling of M.HpaII to one strand is expected to hyperstabilize the underlying DNA duplex. Interestingly, we sometimes observed transient stalling of leading strands at the −38 to −44 positions at a stable, leading strand DPC in mock-depleted egg extracts (e.g. Figure S3F, lane 1). This distal arrest was pronounced in the absence of RTEL1 (Figure 3C, 3D, S3F) and further enhanced by the addition of catalytically inactive RTEL1 (Figure 3C). We propose that −38 to −44 arrest occurs when CMG collides with M.HpaII that is interacting tightly with the underlying double-stranded DNA; the advance from −38 to −30 reflects RTEL1-mediated unwinding of this DNA, which disrupts the M.HpaII nucleoprotein complex, allowing CMG to get closer to the adducted nucleotide (Figure S3G). To further explore this idea, we placed a DPC on the lagging strand template, where it should stabilize the underlying DNA without blocking the translocation strand. As shown in Figure 3D, RTEL1 depletion delayed bypass of the lagging strand DPC, but the delay was less pronounced than the effect seen with a leading strand DPC (Figure 3D and Figure S3F). Interestingly, the arrest at the lagging strand DPC occurred solely at the −35 position, consistent with the absence of a second stalling event of CMG at the adducted nucleotide. Collectively, these observations suggest that RTEL1 not only destabilizes the DNA underlying the DPC but also specifically assists CMG in overcoming leading strand obstacles.

### Efficient DPC proteolysis requires RTEL1

As shown in Figure 1C, the kinetics of CMG bypass were identical whether or not the DPC was degraded. This observation strongly implies that CMG normally bypasses the DPC before proteolysis, and it raises the question of whether bypass is a pre-requisite for proteolysis. To address this point, we replicated a plasmid containing non-methylated leading strand DPCs in undepleted extract so that both proteolysis pathways were active. The extract was either mock-depleted or depleted of RTEL1. At different times, we isolated the plasmid under stringent conditions so that only M.HpaII that was covalently attached to DNA was recovered. After DNA digestion, immunoblotting revealed M.HpaII polyubiquitylation, followed by a decline in M.HpaII levels, which reflects replication-dependent proteolysis (Figure 4A, lanes 1-6; Gao et al., submitted). Strikingly, M.HpaII degradation was greatly delayed in extracts depleted of RTEL1 (Figure 4A, lanes 7-12). This defect was rescued by wild type RTEL1 but not the ATPase mutant (Figure 4A, lanes 13-24; see graph for quantification). If, as the above results indicate, CMG bypass normally precedes DPC proteolysis, then RTEL1 depletion should inhibit approach of the leading strand to the DPC even when proteolysis is allowed. Indeed, approach was delayed in the absence of RTEL1 ATPase activity, consistent with RTEL1 being needed for CMG bypass of the DPC, even when the DPC can be degraded (Figure 4B). In the absence of RTEL1, M.HpaII proteolysis is delayed but not eliminated (Figure 4A-B), and this can be explained by significant CMG bypass still observed in this setting. Together, the data indicate that RTEL1-mediated unwinding past the DPC is essential for its efficient proteolysis.

**Figure 4.**
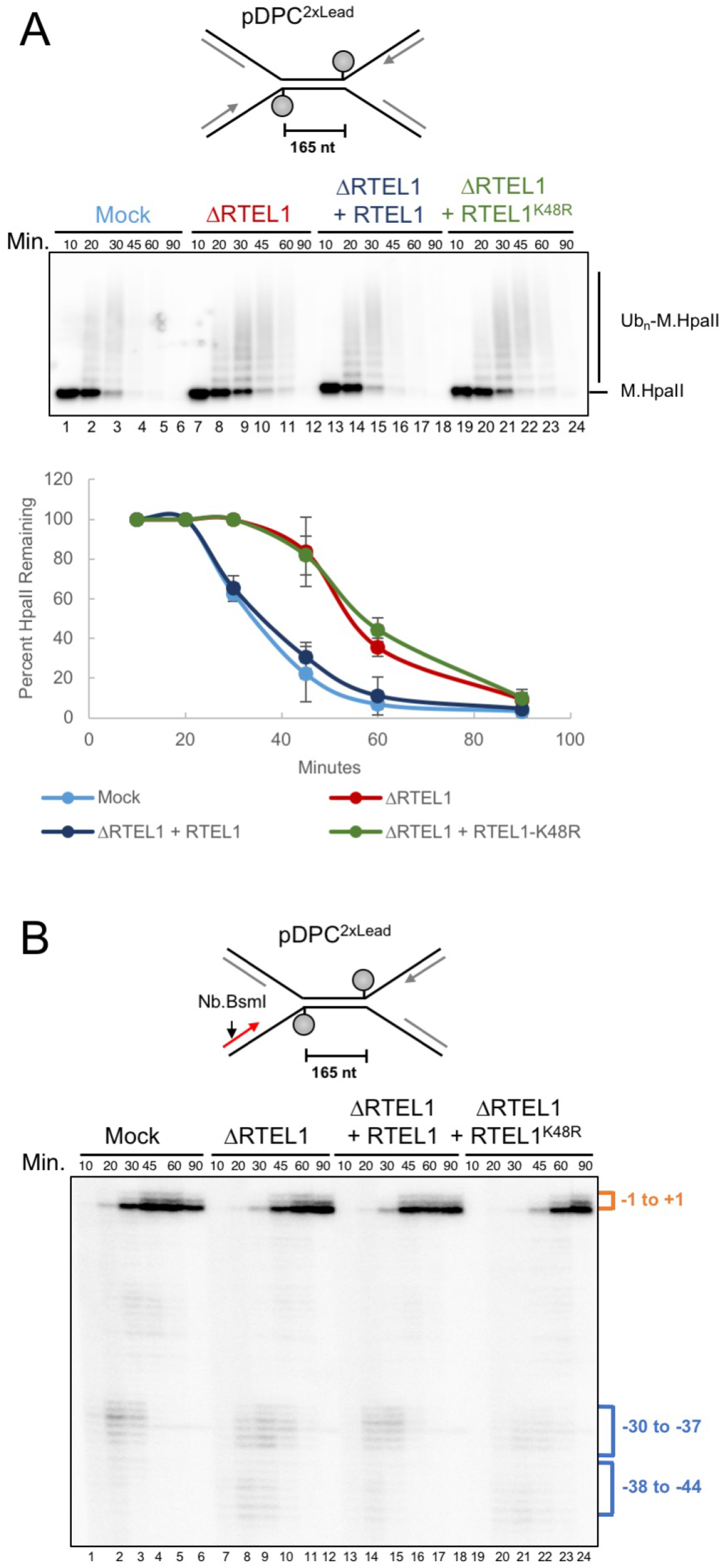
RTEL1 is required for efficient DPC proteolysis. (**A**) pDPC^2xLead^ was replicated in mock-depleted or RTEL1-depleted egg extract and supplemented with buffer, wild-type RTEL1, or RTEL1-K48R. At the indicated times, plasmid was recovered under stringent conditions, the DNA digested, and the resulting samples subjected to immunoblot analysis with M.HpaII antibody. Signal from the entire lane was quantified, and peak signal was assigned a value of 100%. The mean of three independent experiments is shown. Error bars represent standard deviation. (**B**) Parallel reactions to those in (A) were supplemented with [α-^32^P]dATP. Leading strand approach was visualized as in Figure 1G.

### RTEL1-dependent DPC bypass promotes SPRTN activity

As shown in the accompanying manuscript (Gao et al., submitted), DPC repair in egg extracts involves two proteases, SPRTN and the proteasome, which have overlapping functions. To examine whether RTEL1 is required for the SPRTN proteolysis pathway, we examined meDPC, which cannot be ubiquitylated and therefore is not susceptible to the proteasome but can still be degraded by SPRTN (Gao et al., submitted). The action of SPRTN was visible from the appearance of a specific M.HpaII cleavage product that was absent in ΔSPRTN extract (Figure 5A; lanes 1-10; Gao et al., submitted). In the absence of RTEL1, accumulation of the SPRTN-specific product was delayed (Figure 5A; compare lanes 1-5 and 11-15). As seen for DPC bypass, these defects were rescued by wild type but not ATPase deficient RTEL1 (Figure 5A, lanes 16-25), indicating that DNA unwinding past the DPC by RTEL1 is required for SPRTN activity. To address whether CMG bypass itself is required, we examined the effect of tandem leading strand meDPCs since these severely impair CMG bypass (Figure 2A, lanes 13-18), presumably without affecting RTEL1’s ability to unwind past the adducts. As shown in Figure 5B, tandem meDPCs severely inhibited the appearance of the SPRTN-specific M.HpaII product compared to single meDPCs, which allow CMG bypass. Our data indicate that RTEL1-dependent DNA unwinding and CMG bypass of the DPC are both required for the SPRTN pathway.

**Figure 5.**
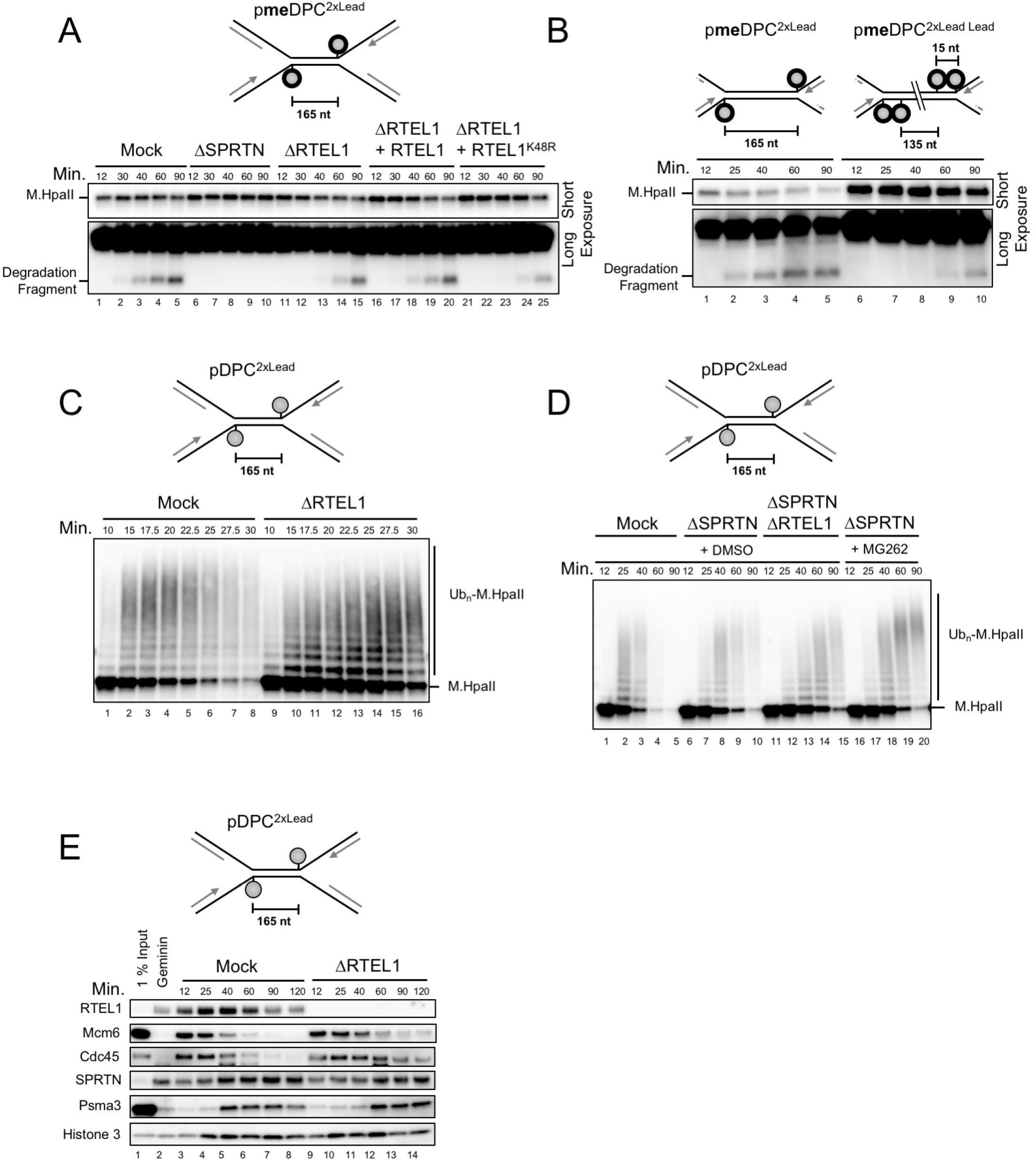
RTEL1 promotes the SPRTN and proteasome pathways. (**A**) pmeDPC^2XLead^ was replicated in mock-depleted, SPRTN-depleted, or RTEL1-depleted egg extract and supplemented with buffer, wild-type RTEL1, or RTEL1-K48R, as indicated. Samples were withdrawn and processed by the stringent pull-down procedure described in Figure 4A. Short and long (larger excerpt) exposures of the same blot are shown. (**B**) pmeDPC^2xLead^ or pmeDPC^2xLead Lead^ were replicated in non-depleted extract. Stringent plasmid pull downs were performed as in Figure 4A and presented as in (A). (**C**) pDPC^2XLead^ was replicated in mock-depleted or RTEL1-depleted egg extracts and stringent plasmid pull-down was performed and analyzed as in Figure 4A. Functional RTEL1 depletion was verified in Figure S4A. (**D**) pDPC^2XLead^ was replicated in mock-depleted, SPRTN-depleted, RTEL1-depleted, or SPRTN/RTEL1-depleted egg extracts that also contained DMSO or MG262. Stringent plasmid pull-down was performed as in Figure 4A. (**E**) pDPC^2XLead^ was replicated in mock-depleted or RTEL1-depleted egg extracts. At different times, plasmid-associated proteins were recovered under non-stringent conditions and blotted with the indicated antibodies.

We next addressed whether RTEL1 affects the proteasome pathway. In Figure 4A (30 minute time point), we noticed a delay in the appearance of highly ubiquitylated M.HpaII species in RTEL1-depleted extracts (lane 3 vs. 9), suggesting that RTEL1 is required for efficient DPC ubiquitylation. A finer time course confirmed this conclusion (Figure 5C), suggesting there might be a defect in proteasome-mediated DPC destruction in the absence of RTEL1. To specifically examine the effect of RTEL1 on the proteasome pathway, we replicated pDPC in SPRTN depleted-extract with or without additional RTEL1 depletion and examined DPC proteolysis, using MG262 addition as a positive control for proteasome inhibition. As shown in Figure 5D, RTEL1 depletion stabilized M.HpaII to a similar extent as MG262 in ΔSPRTN extract (compare lanes 11-15 with 16-20), consistent with RTEL1 functioning in the proteasome pathway. Finally, in RTEL1-depleted extracts, chromatin-binding of SPRTN was reduced, and binding of the PSA3 proteasome subunit was delayed (Figure 5E). Together, our experiments indicate that RTEL1 is required for both proteolysis pathways.

Although the requirement for RTEL1’s ATPase activity in DPC proteolysis strongly argues it functions by unwinding DNA at the DPC, we wanted to address whether it also plays a more direct role, for example by recruiting one or both proteases. To address this possibility, we took advantage of the fact that a ssDNA gap across from the DPC stimulates DPC proteolysis in the absence of a replication fork, obviating the need for CMG bypass or DNA unwinding at the DPC (Figure S4B, lanes 1-6; Gao et al., submitted). Importantly, RTEL1 depletion had no effect on DPC ubiquitylation or proteolysis in this replication-independent setting (Figure S4B, lanes 7-12). Similarly, production of the SPRTN cleavage product from meDPC placed in a ssDNA gap did not depend on RTEL1 (Figure S4C). These data show that in the presence of ssDNA, RTEL1 is not required for SPRTN activity or overall DPC proteolysis. Rather, the primary function of RTEL1 in DPC proteolysis is to unwind DNA around the adduct, which facilitates both proteolysis pathways.

### RTEL1 is required for efficient bypass of a non-covalent nucleoprotein complex

As shown in Figure 3D, RTEL1 is required to efficiently bypass a lagging strand DPC, probably because it helps unwind the DNA underlying the DPC. If this interpretation is correct, it follows that RTEL1 should be required for the replicative bypass of any nucleoprotein complex that stabilize the duplex, even ones that are non-covalent. To address this question, we replicated a plasmid containing an array of 32 *lacO* sites bound by LacR. At different time points, plasmid was recovered and cut with XmnI at a single site before native gel electrophoresis. We previously showed that in this setting, replication forks converge on the outer edges of the LacR array, generating a discrete X-shaped intermediate whose mobility gradually decreases as forks slowly progress through the array (Figure 6A, lanes 1-6; Dewar et al., 2015). When forks meet, the X-shaped species are converted into fully replicated, linear daughter molecules. As shown in Figure 6A, RTEL1 depletion severely delayed the accumulation of linear molecules (lanes 7-12), and this effect was partially rescued by RTEL1^WT^ but not RTEL1^K48R^, which slightly delayed DNA replication (lanes 13-24). To examine fork progression through the LacR array at higher resolution, DNA was nicked near the *lacO* sites with Nt.BspQI, and the radioactive nascent strands were separated on a urea PAGE gel, which reveals fork pausing ~30 nt upstream of each *lacO* site (Figure 6B, lanes 1-6; (Dewar et al., 2015)). Based on this analysis, fork progression through the array was also severely compromised in extracts lacking RTEL1 (Figure 6B, lanes 1-12, red boxes), and the defect was reversed with RTEL1^WT^ but not RTEL1^K48R^ (Figure 6B, lanes 13-24). We conclude that RTEL1 is required for the bypass of covalent and non-covalent nucleoprotein complexes.

**Figure 6.**
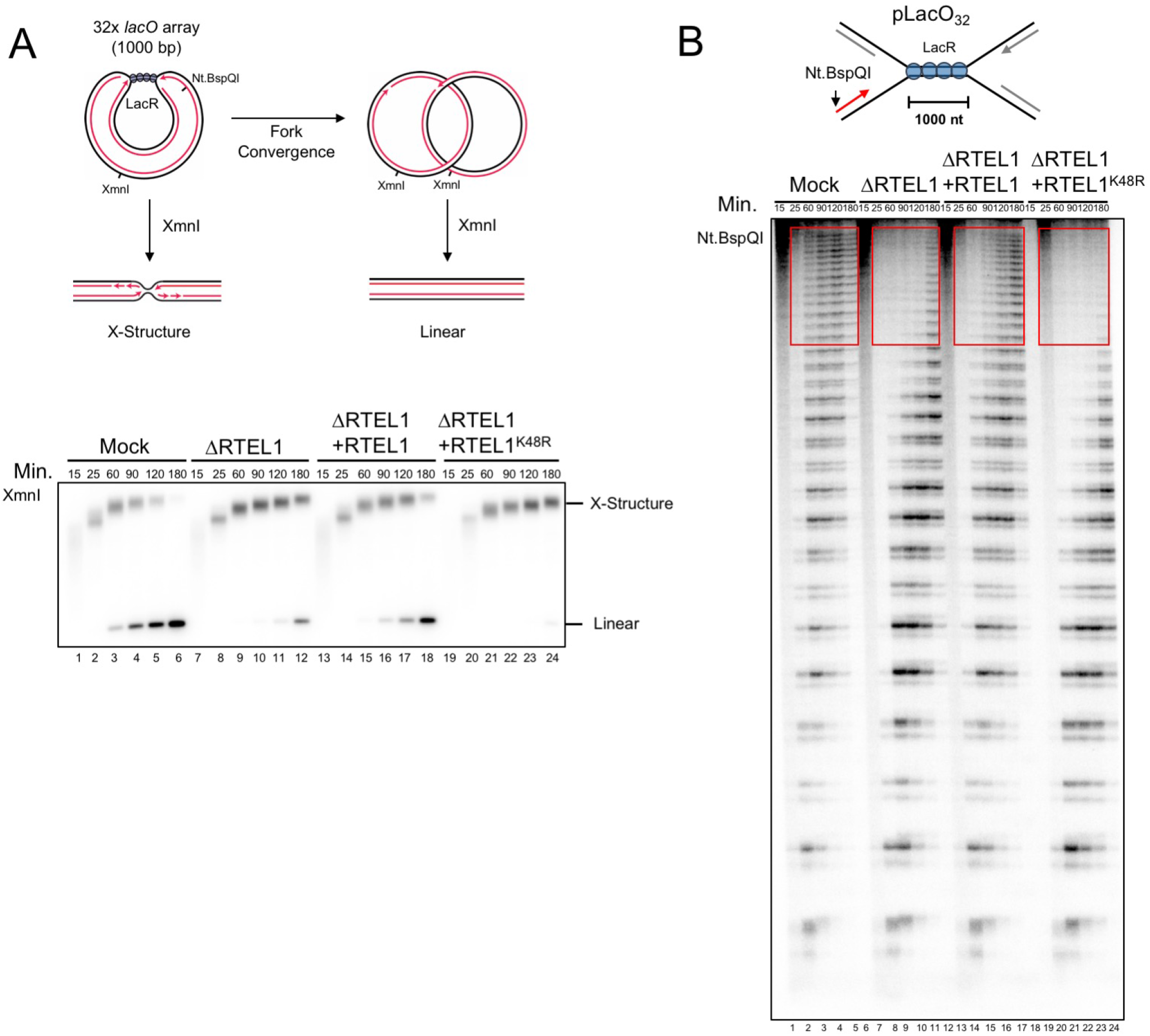
RTEL1 is required for CMG bypass of non-covalent nucleoprotein complexes. (**A**) Top, structures generated with and without XmnI digestion before and after forks pass through the LacR array. pLacO_32_ was pre-incubated with LacR and replicated in mock-depleted or RTEL1-depleted egg extracts containing [α–^32^P]dATP and supplemented with buffer, wild-type RTEL1, or RTEL1-K48R. At the indicated times, DNA was recovered, digested with the single cutter XmnI, resolved by native agarose gel electrophoresis, and visualized by autoradiography. (**B**) DNA samples from (A) were nicked with Nt. BspQI to release the rightward leading strand (red arrow), separated on a denaturing polyacrylamide gel, and visualized by autoradiography.

## Discussion

We present evidence that RTEL1 plays a central role in overcoming nucleoprotein complexes in a vertebrate cell-free system. Surprisingly, RTEL1 endows CMG with the ability to bypass a 45 kDa protein coupled covalently to the translocation strand. RTEL1-dependent CMG bypass normally *precedes* and is required for efficient DPC proteolysis. RTEL1 also promotes fork progression past transcription factors bound non-covalently to DNA. Our results establish a comprehensive framework to understand how the vertebrate replisome overcomes covalent and non-covalent nucleoprotein obstacles (Figure 7).

**Figure 7.**
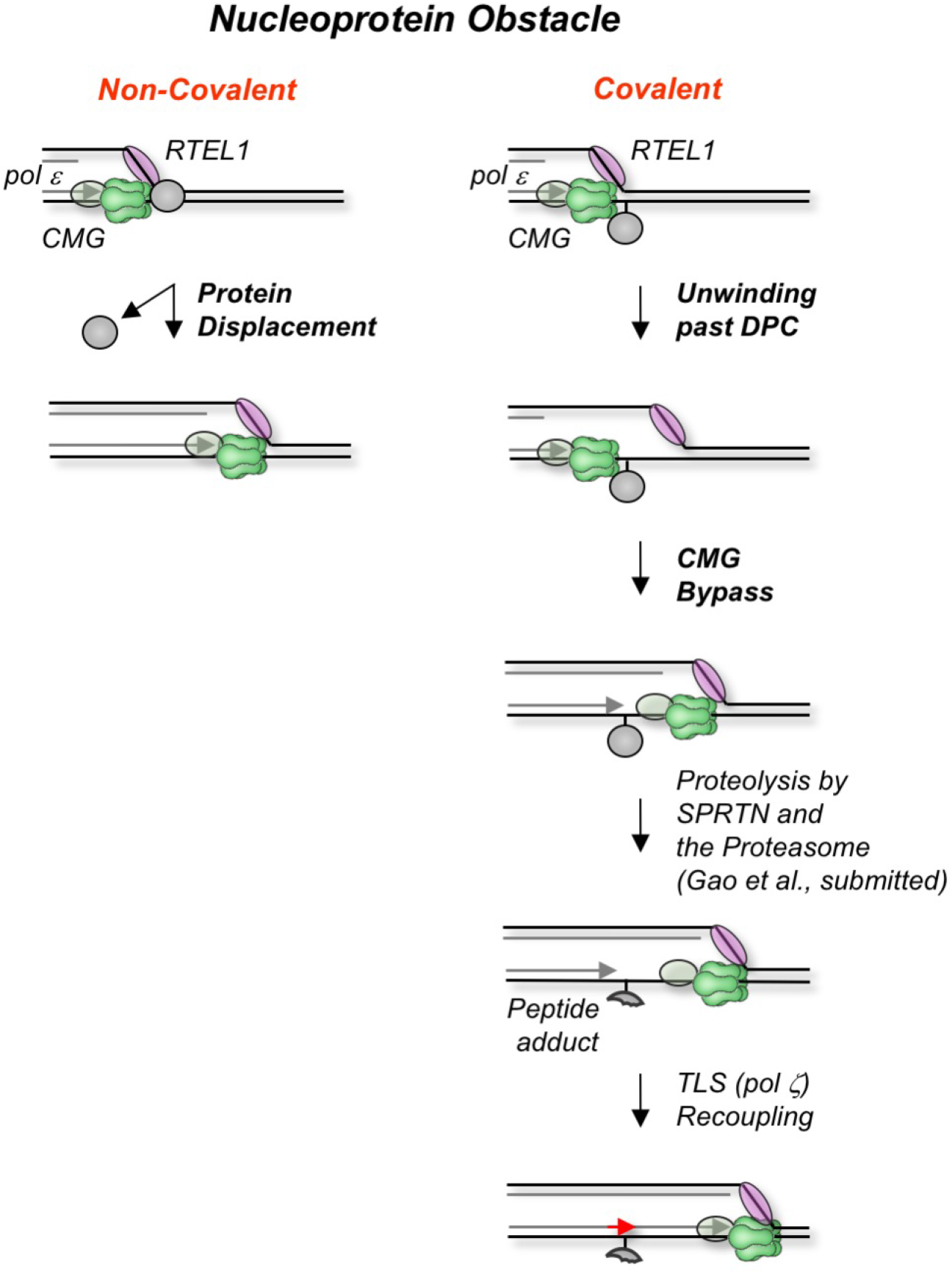
Model for RTEL1 bypass of nucleoprotein complexes. When the replisome encounters a nucleoprotein barrier, CMG stalls and RTEL1 is either recruited de novo or, if it normally travels with the fork as suggested by our data, it becomes engaged. In the case of non-covalent nucleoprotein complexes, RTEL1 and CMG cooperate to unwind the DNA underlying the protein, leading to its displacement and immediate resumption of fork progression. At a covalent DPC, RTEL1 translocates along the undamaged lagging strand template, exposing ssDNA at the DPC and beyond. The generation of ssDNA facilitates CMG bypass of the DPC. Given the high affinity of pol ε (grey oval) for CMG, we envision that it bypasses the DPC with CMG. Once CMG has bypassed the DPC, the DPC undergoes proteolysis by SPRTN or the proteasome (see Gao et al., submitted). Finally, the leading strand is extended past the peptide adduct using translesion synthesis polymerases.

### Disruption of non-covalent nucleoprotein complexes by RTEL1

The most common nucleoprotein complexes encountered by replication forks are likely to be non-covalent in nature. In bacteria, where the replicative DnaB helicase travels 5’ to 3’ on the lagging strand template, bypass of such complexes involves the function of the 3’ to 5’ DNA helicases Rep or UvrD, which operate on the leading strand template. In yeast cells, the situation is reversed: bypass involves Rrm3, whose 5’ to 3’ polarity places it on the lagging strand template, opposite CMG. We showed that CMG progression past a LacR array in a vertebrate model system requires RTEL1, which translocates on the lagging strand template. A conserved mechanism emerges in which replication forks employ an accessory helicase on the strand opposite the one hosting the replicative DNA helicase.

In egg extracts and cells, RTEL1 is constitutively associated with replication forks in the absence of experimentally-induced DNA damage (Figure 3A; Vannier et al., 2013), suggesting RTEL1 might travel with the replisome. We previously characterized a vertebrate replication progression complex (RPC) that is retained on chromatin when CMG unloading during DNA replication termination is blocked. The RPC contains, among other things, CMG, DNA pol ε, MCM10, CLASPIN, TIMELESS-TIPIN, and AND-1 (yeast Ctf4)(Dewar et al., 2017). Notably, RTEL1 was the only 5’ to 3’ DNA helicase detected in the RPC (See Figures 2B and S4D in (Dewar et al., 2017)). The interaction of RTEL1 with the replisome positions this helicase for rapid engagement when obstacles are encountered but might also allow RTEL1 and CMG to cooperate in tackling barriers. Whatever the precise mechanism, we propose that RTEL1 disrupts non-covalent nucleoprotein complexes by unwinding the underlying DNA, which will, in most cases, displace the protein obstacle from DNA.

### Bypass of a DPC on CMG’s translocation strand

Less commonly, forks will encounter covalent DPCs. We previously showed that the disappearance of the CMG footprint at a leading strand DPC (approach) correlates with the latter’s proteolysis, and that when DPC proteolysis is blocked due to ubiquitin depletion, loss of the footprint is dramatically delayed. On this basis, we proposed that DPCs block CMG progression (Duxin et al., 2014). Contrary to this idea, we now present evidence that CMG can readily bypass a stable DPC on CMG’s translocation strand. First, when DPC proteolysis is blocked without ubiquitin depletion (by inhibiting the SPRTN and proteasome pathways), the leading strand approaches the DPC in the presence of p97-i, which blocks CMG unloading. From this observation, it can be inferred that the CMG has to bypass the DPC to allow approach. Second, CMG is detected at an ICL downstream of an intact DPC, consistent with a bypass mechanism. Although we have not formally demonstrated that the same molecule of CMG that collides with the DPC also undergoes bypass, such a mechanism is highly likely because in the absence of CMG unloading, the only plausible mechanism by which the stalled CMG could get out of the way of the advancing leading strand is via bypass. In addition, *de novo* MCM2-7 complex recruitment to chromatin in S phase has never been observed, making it unlikely that a new CMG could be recruited on the other side of the DPC. Finally, we previously showed that purified large T-antigen can bypass M.HpaII on the translocation strand, establishing a precedent for DPC bypass (Yardimci et al., 2012). These considerations lead to the remarkable conclusion that CMG overcomes large adducts on its translocation strand. Our previous observation that approach was inhibited when DPC proteolysis was blocked by ubiquitin depletion was most likely due to pleiotropic consequences of this manipulation (Duxin et al., 2014).

We envision two possible mechanisms of bypass. In the first, the MCM2-7 ring opens, allowing CMG to slide past the DPC (Figure S5A). During licensing, MCM2-7 opens at the MCM2-MCM5 interface to allow entry of dsDNA into the MCM2-7 central channel (Bochman and Schwacha, 2010; Samel et al., 2014). Moreover, during initiation, one strand of dsDNA is probably extruded from MCM2-7 through an unknown mechanism. Therefore, MCM2-7 is highly dynamic, and it is conceivable that ring-opening could underlie DPC bypass. However, a large gap in the ring might have to be created to accommodate an adduct as large as M.HpaII (45 kDa), and whether MCM2-7 is flexible enough to accomplish this is uncertain. Use of the MCM2-MCM5 gate for DPC bypass would likely require transient dissociation of Cdc45 and/or GINS (Hashimoto et al., 2011), which both protect the gate in CMG. In a second possible mechanism, CMG threads the denatured DPC through its central channel (Figure S5B). Because most DPCs will be attached to DNA at an internal amino acid, this mechanism would require CMG’s central pore to accommodate ssDNA and two polypeptide chains. Future manipulation of CMG and the DPC will be required to distinguish between these models. CMG bypass appears to be distinct from traverse, when a replication fork moves past an intact ICL (Huang et al., 2013), since traverse but not bypass requires FANCM and ATR ((Huang et al., 2013; Ling et al., 2016); our data not shown)).

It is interesting to consider how RTEL1 might promote bypass. As shown in Figure S3E, the requirement for RTEL1 in CMG bypass is largely abolished when a ssDNA bubble is placed downstream of the DPC. This observation suggests that the primary function of RTEL1 is to generate ssDNA downstream of the lesion, and it is difficult to reconcile with a model in which CMG denatures the DPC (Figure S5B), as denaturation should not be facilitated by ssDNA downstream. We favor the idea that ssDNA created beyond the DPC allows the breached CMG to re-engage with DNA downstream of the DPC. Given recent evidence that the non-catalytic N-terminal tier of CMG resides at the leading edge of the fork (Douglas et al., 2018; Georgescu et al., 2017), the N-terminus might re-close around ssDNA while the c-terminal ATPase domain has still not bypassed the DPC. According to the recently proposed “inchworm” mechanism of CMG unwinding, the N-terminal tier of CMG might reach past the DPC when CMG is in the extended conformation (Yuan et al., 2016). The evidence discussed above that RTEL1 resides in a large complex containing CMG suggests that RTEL1 might help “tow” CMG past the DPC, although this does not appear to be essential. Future structural and biochemical analysis will be required to define the mechanism of RTEL1-mediated CMG bypass.

### CMG bypass precedes and is required for efficient DPC proteolysis

Strikingly, our data indicate that CMG normally bypasses the DPC *before* proteolysis and that CMG bypass is required for efficient proteolysis. First, we see no difference in the kinetics of leading strand approach when proteolysis proceeds unimpeded compared to when it is completely inhibited (Figure 1C). This result strongly implies that CMG bypass normally precedes proteolysis. Second, when we deplete RTEL1, which inhibits CMG bypass, proteolysis in general and SPRTN activity in particular are severely affected, and the defects are not rescued by ATPase deficient RTEL1. Third, tandem leading strand DPCs, which inhibit bypass in the presence of RTEL1, also severely inhibit SPRTN activity (Figure 5B). Finally, DPC proteolysis by SPRTN requires that the leading strand advance to within a few nucleotides of the DPC (Gao et al., submitted), which is only possible if CMG has moved out of the way. Altogether, the data clearly support the idea that CMG can bypass an intact DPC, and that such bypass is a pre-requisite for DPC proteolysis by SPRTN (Figure 7).

While the evidence discussed above strongly suggests that CMG bypass is required for SPRTN activity, the relationship between bypass and the proteasome is less clear. RTEL1 depletion appears to impair the production of long ubiquitin chains on the DPC, and it phenocopies MG262 addition in SPRTN-depleted extracts, clearly implicating RTEL1 in the proteasome pathway. However, it is unclear whether RTEL1-dependent DNA unwinding at the DPC is sufficient to trigger proteasome activity, or whether CMG bypass is also required. Consistent with the former possibility, Gao et al. (submitted) showed that placing a DPC in a ssDNA context is sufficient to trigger proteasome-mediated DPC proteolysis and that leading strand approach to the lesion is not required. A mechanism in which proteasome activity requires RTEL1 unwinding but not CMG bypass would allow the destruction of DPCs that cannot be bypassed (e.g. because they are too large). In this scenario, DPCs are normally bypassed by CMG before proteolysis by SPRTN and the proteasome (Figure 7), but in the case of impassable DPCs, proteolysis by the proteasome would happen first, providing a backup mechanism to prevent CMG stalling (not shown).

The “bypass first” mechanism we describe has features that could enhance genome maintenance. First, moving CMG past the DPC before proteolysis reduces the probability that the helicase is accidentally destroyed. Because there is no known pathway to reload CMG from scratch in S phase, avoiding inadvertent CMG inactivation is crucial to prevent fork collapse. In contrast, if DNA polymerases or other replisome components should be degraded, they can readily rebind. Second, if CMG bypass occurs but TLS fails, the lagging strand is still extended past a leading strand DPC (Figure 1C; upper autoradiogram). In this case, the leading strand could be extended past the DPC via template switching (Figure S5C) or repriming. Other implications of the “bypass first” mechanism will likely emerge from future studies.

Mammalian RTEL1 exhibits properties consistent with the idea that it helps overcome nucleoprotein complexes. While homozygous RTEL1 mutations are lethal, deletion of RTEL1’s PIP box, with which it binds to PCNA, leads to defective replication fork progression in the absence of external insults, suggesting that RTEL1 helps overcome endogenous barriers to replication fork progression (Vannier et al., 2013). In egg extracts, the PIP box of RTEL1 is not required to bypass DPCs (data not shown), indicating that in this setting, RTEL1 can bind to replication forks via other means. We found that even when RTEL1 is depleted by 95%, there is still significant DPC bypass and proteolysis. Although we cannot rule out that residual RTEL1 mediates these effects, we suspect there could be redundancy among the several vertebrate 5’ to 3’ helicases. At present, it is unclear whether any of the phenotypes observed in RTEL1-deficient mice or humans are attributable to defective CMG bypass of DPCs or other obstacles.

## Conclusion

Our data suggest a unified model for the interaction of eukaryotic replication forks with nucleoprotein complexes (Figure 7). When CMG stalls at a barrier, RTEL1, which appears to travel with the fork, unwinds DNA towards the obstacle. For non-covalent complexes, the combined action of RTEL1 and CMG unwinds the DNA underlying the obstacle, leading to displacement of the protein and immediate resumption of CMG movement. For covalent complexes, RTEL1 unwinds DNA past the DPC, leading to CMG bypass of the DPC and activation of proteolysis.

## Acknowledgements

We thank Niels Mailand, Peter Burgers, and members of the Walter laboratory for feedback on the manuscript. J.L.S. is supported by a Damon Runyon Fellowship. J.C.W. is supported by NIH grant HL98316. J.C.W. is an investigator of the Howard Hughes Medical Institute.

## Author Contributions

J.L.S. and J.C.W. designed and analyzed the experiments and prepared the manuscript. J.L.S. performed the experiments. A.G. developed the protocol to methylate DPCs. M.R. and M.M. performed and analyzed the mass spectrometry. J.P.D. and N.B.L. developed and supplied the pDPC^ssDNA^ and pmeDPC^ssDNA^ plasmids. J.P.D. developed the stringent plasmid pull-down protocol.

## Methods

All experiments were performed three or more times, except for Figures 3D, S4B and S4C, which were performed twice.

### Preparation of DNA Constructs

To generate pDPC, we first created pJLS2 by replacing the AatII-BsmBI fragment from pJD2 (Duxin et al., 2014) with the a sequence (5’-GGGAGCTGAATGCCGCGCGAATAATGGTTTCTTAGACGT-3’) which contains a Nb.BsmI site. To generate pDPC^2xLead^ the SacI-BssHII fragment from pJLS2 was replaced with the following sequence: 5’-CATCCACTAGCCAATTTATGCTGAGGTACCGGATTGAGTAGCTACCGGATGCTGAGGGGATCCACTAGCCAATTTATCATGG-3’. To generate pDPC^ICL^, we first created pJLS6 by replacing the SacI-BssHII fragment of pJLS2 with a sequence (5’-CGAAGACAACCCTTAAACAGCTGAAAGAAGACGCGTGCG-3’) that contains two BbsI sites. The cisplatin ICL containing oligonucleotides were inserted as previously described (Enoiu et al., 2012). To generate pDPC^Lag/Lead^, and pDPC^Lag/Lead-Bubble^ we first replaced the BssHII-KpnI fragment from pJLS2 with the sequence 5’-CGCGCTTAATCAGTGAGGCACCTATCTCCGGTCTGAGTCATGCGTAACTCGAGTGCTTGTAGTGGATTTACCGGATTGAGTAGCTACCGGATGGTAC-3’ hybridized with either 5’-CATCCGGTAGCTACTCAATCCGGTAAATCCACTACAAGCACTCGAGTTACGCATGACTCAGACCGGAGATAGGTGCCTCACTGATTAAG-3’ or CATCCGGTAGCTACTCAATCCGGTATTAGGTGATGTTCGTGAGCTCAATGCGTACTGAGTCTGGCGAGATAGGTGCCTCACTGATTAAG-3’, respectively. The supercoiled band was purified by cesium chloride gradient ultracentrifugation and DNA was subsequently crosslinked to methylated M.HpaII-His_6_, as previously described (Duxin et al., 2014). pJLS2, pJLS6^ICL^, or pJLS3 were nicked with Nt.BbvcI and ligated with an oligonucleotide containing a fluorinated cytosine (5’-TCAGCATCCGGTAGCTACTCAATC[C5-Fluor dC]GGTACC-3’) and subsequently crosslinked to M.HpaII-His_6_ or methylated M.HpaII-His_6_ to generate pDPC^Lead^, pDPC^ICL^, pDPC^2xLead^ or pmeDPC^Lead^, pmeDPC^ICL^, and pmeDPC^2xLead^, respectively, as previously described (Duxin et al., 2014). To create pmeDPC^Lead/Lag^, pJLS2 was first nicked with Nt.BbvcI and ligated with an oligonucleotide containing a fluorinated cytosine (5’-TCAGCATC[C5-Fluor dC]GGTAGCTACTCAATCCGGTACC-3’). It was subsequently nicked with Nb.BbvcI and ligated with a second oligonucleotide containing a fluorinated cytosine (5’-TGAGGTAC[C5-Fluor dC]GGATTGAGTAGCTACCGGATGC-3’) before crosslinking to methylated M.HpaII-His_6_. To create pmeDPC^Lag/Lead^ pJLS2 was first nicked with Nt.BbvcI and ligated with an oligonucleotide containing a fluorinated cytosine (5’-TCAGCATCCGGTAGCTACTCAATC[C5-Fluor dC]GGTACC-3’). It was subsequently nicked with Nb.BbvcI and ligated with a second oligonucleotide containing a fluorinated cytosine (5’-TGAGGTACCGGATTGAGTAGCTAC[C5-Fluor dC]GGATGC-3’) before crosslinking to methylated M.HpaII-His_6_. To create pmeDPC^Lead/Lead^ or pmeDPC^2xLeadLead^, pJLS2 or pJLS3 was nicked with Nt.BbvcI and ligated with an oligonucleotide containing two fluorinated cytosines (5’-TCAGCATC[C5-Fluro dC]GGTAGCTACTCAATC[C5-Fluro dC]GGTACC-3’) and subsequently crosslinked to methylated M.HpaII-His_6_ to generate pmeDPC^Lead/Lead^ or pDPC^2xLeadLead^, respectively, as previously described (Duxin et al., 2014). Creation of pDPC^ssDNA^ and pmeDPC^ssDNA^ is described in (Gao et al. submitted). Creation of pLacO_32_ was previously described (Dewar et al., 2015).

### *Xenopus* Egg Extracts and DNA Replication

*Xenopus* egg extracts were prepared essentially as described (Lebofsky et al., 2009). Briefly, licensing was carried out by supplementing a high-speed supernatant (HSS) of egg cytoplasm with plasmid DNA at a final concentration of 7.5–15 ng/μL. For radiolabeling DNA replication products, [α-^32^P] dATP was added to HSS prior to the DNA. For replication in the presence of LacI, 1 volume of plasmid (75 ng/μL) was incubated with an equal volume of 12 μM LacI for 30 minutes prior to transfer into HSS so that the final concentration of plasmid was 7.5 ng/μl (Duxin et al.). Licensing mixes were incubated for 30 min at room temperature to assemble pre-replicative complexes (pre-RCs). To prevent licensing, Geminin was added to HSS at a final concentration of 10 μM and incubated for 10 min at room temperature prior to addition of plasmid DNA. To initiate replication, 1 volume of licensing reaction was mixed with 2 volumes of nucleoplasmic extract (NPE) that had been diluted two-fold with 1xELB-sucrose (10 mM Hepes-KOH pH 7.7, 2.5 mM MgCl_2_, 50 mM KCl, 250 mM sucrose). 0.5 μl aliquots of replication reaction were typically stopped with 5–10 volumes of replication stop buffer (8 mM EDTA, 0.13% phosphoric acid, 10% ficoll, 5% SDS, 0.2% bromophenol blue, 80 mM Tris-HCl at pH 8), treated with 1 μg/μL Proteinase K. For nascent strand analysis, 2.5 μl aliquots of replication reaction were stopped in 10 volumes of sequencing stop buffer (0.5 % SDS, 25 mM EDTA, 50 mM Tris-HCl pH 8.0) followed by addition of 1.25 μl of 190 ng/μL RNase A and incubated for 30 minutes at 37 °C. After RNase digestion, 1.25 μl of 900 ng/μL Proteinase K was added to the DNA samples and incubated overnight at room temperature. Following the Proteinase K treatment, samples were diluted to 150 μl with 10 mM Tris-HCl pH 8.0. The samples were extracted once with an equal volume of phenol/chloroform followed by one extraction with an equal volume of chloroform. The DNA was then precipitated with the addition of 0.1 volumes 3M sodium acetate pH 5.2 and 1 μl glycogen (20 mg/ml stock) and resuspended in 7.5 μl. For RTEL1 immunodepletion and rescue experiments, NPE was supplemented with ~ 200 nM recombinant wild type or mutant *Xenopus* RTEL1 and incubated for 15 minutes prior to replication initiation. For Ub-VS treatment, NPE was supplemented with 22.5 μM ubiquitin vinyl sulfone (Ub-VS) (stock ~250 μM; Boston Biochem.) and incubated for 15 minutes prior to mixing with HSS (15 μM final concentration in replication mix). For p97i (NMS873; Sigma) treatment, NPE was supplemented with 200 μM NMS-873 (20 mM stock) and incubated for 10 minutes prior to mixing with HSS (133.33 μM final concentration in replication mix. For MG262 (stock 20 mM; Boston Biochem.) treatment, NPE was supplement with 200 μM MG262 and incubated for 15 minutes prior to mixing with HSS (133.33 μM final concentration in replication mix). A 100 mM Cdc7-i (PHA-767491; Sigma) stock was prepared in distilled water and added to replication mix at a final working concentration of 100 μM at the specified time point. Samples were analyzed by native 0.8% agarose gel electrophoresis. Gels were exposed to phosphorscreens and imaged on a Typhoon FLA 7000 phosphorimager (GE Healthcare). In Figures 1G (top panel) and Figure S1F, the original images were converted into a log scale for display by applying the function f(p) = log(p)*255/log(255) to each pixel (p) in the images. Band or total lane intensities were quantified using Multi-Gauge software (Fujifilm) with subtraction of appropriate background.

### Nascent strand analysis

To digest radio-labeled nascent leading-strands, 3-4 μl of extracted and ethanol precipitated DNA (see above) at 1-2 ng μl^−1^ was incubated in buffer 3.1 (New England BioLabs) with 0.45 units μl^−1^ Nb.BsmI (New England BioLabs) in a 5 μl reaction at 65 °C for 1 h. To nick rightward leading strands of pLacO_32_, 3-4 μl of purified DNA at 1-2 ng μl^−1^ was incubated in buffer 3.1 with 0.4 units μl^−1^ Nt.BspQI (New England BioLabs) at 37 °C for 1 h. Nicking reactions were stopped with 0.5 volumes of Sequencing Stop solution (95% formamide, 20 mM EDTA, 0.05 *%* bromophenol blue, 0.05% xylene cyanol FF). Nicked DNA (3.5 to 4 μl samples) was separated on 4 % (for pLacO) or 7% (pDPC) polyacrylamide sequencing gels. Gels were dried and subjected to phosphorimaging using a Typhoon FLA 7000 phosphoimager. Gels were quantified using Multi Gauge software (Fuji Photo Film Co.). For Figure 6A, 1 μl of purified DNA was used for XmnI digestion.

To quantify the percentage of CMG that underwent bypass in Figures S1E, S2, 2B, 3C, 3D, S3E (called “approach” in Figure S1E, where bypass had not yet been established), the radioactive signal of leading strands located between positions +1 and −28 on the gel (reflecting CMGs that have bypassed) was divided by the radioactive signal for leading strands between positions +1 and −38 (reflecting CMGs that have stalled at the lesion or undergone bypass). In the case of pmeDPC^Lead/Lead^ (Figure 2B, lanes 13-18), we divided the signal between +1 of the 2^nd^ DPC and −1 of the 1^st^ DPC (both DPCs bypassed) by the signal between +1 of the 2^nd^ DPC and −38 of the first DPC (bypassed and not bypassed).

### Antibodies and Immunodepletion

The xlRTEL1-N antibody was raised against a fragment of *Xenopus laevis* RTEL1 encompassing amino acids 400-654, which was tagged on its N-terminus with His_6_. The protein fragment was overexpressed and purified from bacteria under denaturing conditions, and the antibody was raised by Pocono Rabbit Farm & Laboratory. The RTEL1 antibody was affinity purified from the serum using the RTEL1 antigen according to standard protocols. The xlRTEL1-C antibody was raised against amino acids 428-443 (Ac-HPDTSQRKKPRGDIWSC-amide) by New England Peptide. The following antibodies were described previously: CDT1 (Arias and Walter, 2005), Orc2 (Walter and Newport, 1997), Cdc45 (Mimura and Takisawa, 1998), M.HpaII (Gao, et al., submitted), PSA3 (Gao, et al., submitted), SPRTN (Gao, et al., submitted), and Histone H3 (Cell Signaling Cat #9715S). Peptide antibodies against Mcm6 will be described elsewhere.

For RTEL1 immunodepletion, 3.5 volumes of purified RTEL1 antibody (1 mg mL^−1^) or an equivalent amount of rabbit IgG purified from non-immunized rabbit serum (Sigma) were incubated with 1 volume of Protein A Sepharose Fast Flow (PAS) (GE Healthcare) overnight at 4°C. For SPRTN immunodepletion, 4 volumes of SPRTN serum was incubated with 1 volume of Protein A Sepharose Fast Flow (PAS) (GE Healthcare) overnight at 4°C. For mock depletion, 4 volumes of preimmune serum from matched rabbit, was used. One volume of antibody-conjugated Sepharose was then added to 5 volumes of precleared HSS or NPE and incubated for 1 hour at 4°C. The HSS or NPE was collected and incubated two more times with antibody-conjugated sepharose for a total of three rounds of depletion. The depleted HSS or NPE was collected and used immediately for DNA replication, as described above.

### Protein Expression and Purification

M.HpaII-His_6_, LacI-biotin, and LacI-His_6_ were expressed and purified as previously described (Duxin et al. 2014). Lysine methylation of M.HpaII was carried out as previously described (Gao, et al., submitted). *Xenopus* RTEL1 open reading frame with an N-terminal GST tag separated by a 3C cleavage site was cloned into pFastBac1 (Thermo Fisher Scientific) using custom gene synthesis from Integrated DNA Technologies (IDT). The RTEL1 sequence was confirmed by Sanger sequencing. Mutants of RTEL1 were created by around-the-horn site-directed mutagenesis, and mutations were confirmed by Sanger sequencing. The GST-RTEL1 Baculoviruses were made using the Bac-to-Bac system (Thermo Fisher Scientific) according to the manufacturer’s protocols. GST-RTEL1 and mutants were expressed in 3 L suspension cultures of Sf9 cells (Thermo Fisher Scientific) by infection with RTEL1 baculovirus for 36-48 hrs. Sf9 cells were collected via centrifugation and washed with 1XPBS and subsequently pelleted by centrifugation and flash frozen. Cell pellets were thawed and resuspended in an equal volume of 2X Lysis Buffer (100 nM HEPES pH7.5, 1 M NaO_2_Ac, 20 % sucrose, 0.2 % IGEPAL, 4 mM DTT, 2X Roche EDTA-free Complete protease inhibitor cocktail), 1X Lysis Buffer (50 mM HEPES pH7.5, 500 mM NaO_2_Ac, 10 % sucrose, 0.1 % IGEPAL, 2 mM DTT, 1X Roche EDTA-free Complete protease inhibitor cocktail) to the weight of the cell pellet. Cells were lysed by two rounds of sonication, followed by addition of ammonium sulfate (4M stock) to 200 mM final concentration and 45 μl/ml Polymin P (10 % stock) and stirred at 4 °C for 10 minutes. Lysate was cleared by ultracentrifugation at 25,000 rpm in a Beckman Ti45 rotor for 1 hour. The supernatant was subjected to ammonium sulfate precipitation using 0.2 g/ml ammonium sulfate. Proteins were pelleted by ultracentrifugation at 25,000 rpm in a Beckman Ti45 rotor for 1 hour. The supernatant was discarded and protein pellets were resuspended in 50 ml Wash Buffer A_500_ (25 mM HEPES pH7.5, 500 mM NaO_2_Ac, 10 % sucrose, 0.01 % IGEPAL, 2 mM DTT, 1X Roche EDTA-free Complete protease inhibitor cocktails). The resuspended pellet was incubated for 2 hours with 300 μl of Glutatione sepharose™ 4B (GE) at 4 °C. Following incubation, resin was first washed with 20 ml of Wash Buffer A500 and then with 10 ml of Wash Buffer A_200_ (25 mM HEPES pH7.5, 200 mM NaO_2_Ac, 10 % sucrose, 0.01 % IGEPAL, 2 mM DTT, 1X Roche EDTA-free Complete protease inhibitor cocktails). Proteins were eluted from the resin with Elution Buffer E_200_ (25 mM HEPES pH7.5, 200 mM NaO_2_Ac, 10 % sucrose, 0.005 % IGEPAL, 2 mM DTT, 20 mM L-glutathione reduced, pH adjusted to 8.0). Fractions were pooled and dialyzed against Dialysis Buffer (25 mM HEPES pH7.5, 200 mM NaO_2_Ac, 10 % sucrose, 0.005 % IGEPAL, 2 mM DTT) with addition of HRV 3C protease (Thermo Fisher) at 4°C for 4 hr. Aliquots of RTEL1 were flash frozen and stored at −80°C.

### Plasmid Pull-Down

The plasmid pull-down assay was performed as described (Budzowska et al., 2015). Briefly, streptavidin-coupled magnetic beads (Invitrogen; 10 μl per pull-down) were washed three times with 50 mM Tris (pH 7.5), 150 mM NaCl, 1 mM EDTA pH 8, 0.02% Tween-20. Biotinylated LacI was added to the beads (4 pmol per 10 μl beads) and incubated at room temperature for 40 min. The beads were then washed four times with Pull-down Buffer (10 mM HEPES (pH 7.7), 50 mM KCl, 2.5 mM MgCl_2_, 250 mM sucrose, 0.25 mg/ml BSA, 0.02% Tween-20) and resuspended in 40 μl of the same buffer. The bead suspension was stored on ice until needed. At the indicated times, 4.0 μl samples of the replication reaction were withdrawn and gently mixed with LacI-coated streptavidin Dynabeads. The suspension was immediately placed on a rotating wheel and incubated for 30 min at 4 °C. The beads and associated proteins were isolated by centrifugation through a sucrose cushion (10 mM HEPES pH 7.7, 2.5 mM MgCl_2_, 50 mM KCl, 0.5 M sucrose, 0.02 % Tween), then washed once with Pull-down Buffer. All residual buffer was removed, and the beads were resuspended in 20 μl of 2X Laemmli sample buffer. Equal volumes of the protein samples were blotted with the indicated antibodies.

### DPC pull-down

The DPC pull-down assay to quantify how much M.HpaII was removed from the plasmid during DPC repair was performed as described (Gao et al., submitted).

## Supplemental Figure Legends

**Figure S1.**
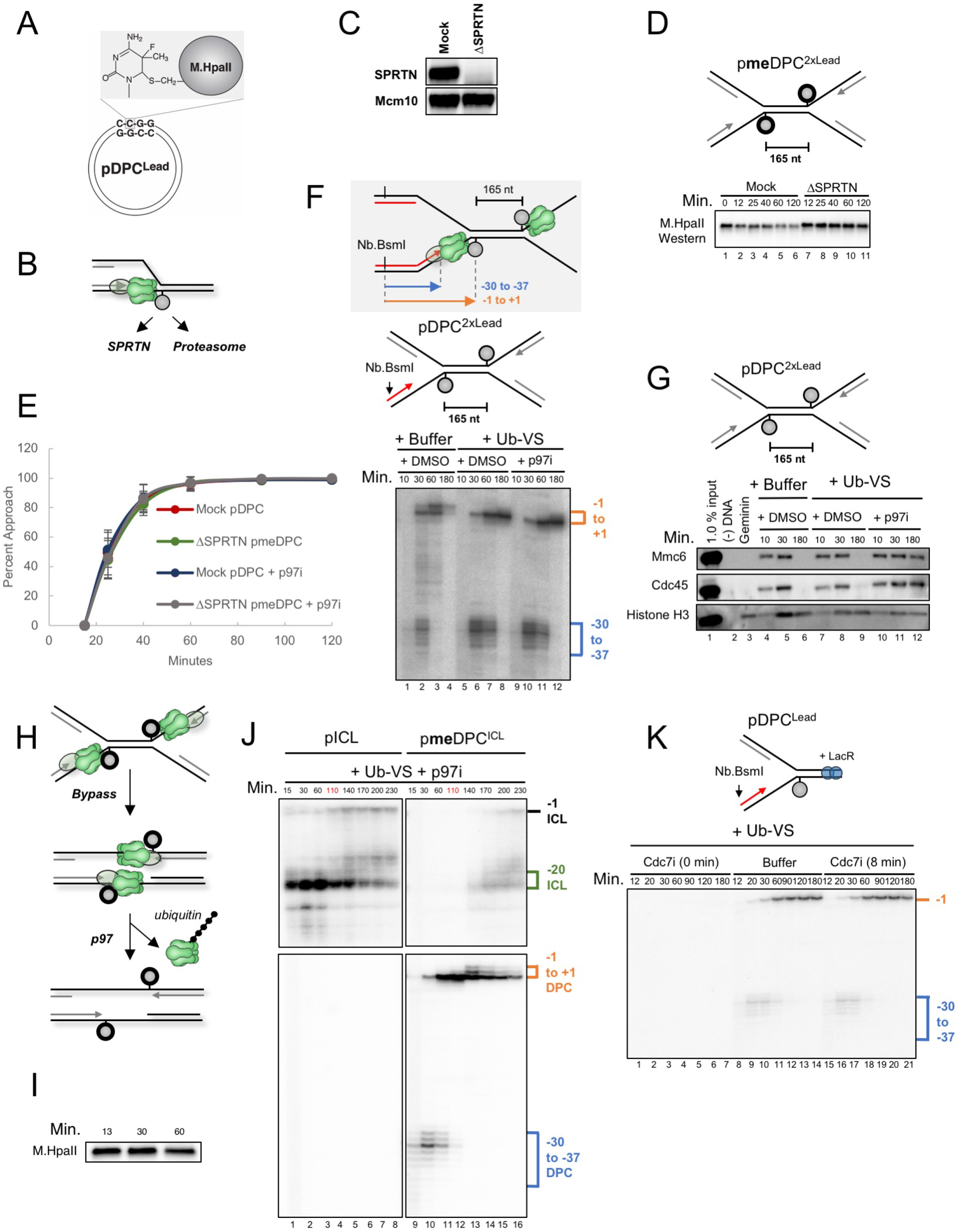
Related to Figure 1. (**A**) Schematic showing trapping of M.HpaII on a plasmid containing a fluorinated M.HpaII recognition site. (**B**) Schematic of the two replication-coupled DPC proteolysis pathways in *Xenopus* egg extracts (Gao et al., submitted). (**C**) Mock-depleted and SPRTN-depleted egg extracts were blotted using SPRTN and Mcm10 (loading control) antibodies. (**D**) pmeDPC^2xLead^ was replicated in the indicated egg extracts. At different times, plasmid was recovered under stringent conditions, the DNA digested, and the released proteins subjected to immunoblot analysis with M.HpaII antibody. (**E**) The percentage of leading strands that underwent approach was quantified based on disappearance of the CMG footprint (see methods), and the mean of five experiments is graphed. Error bars represent the standard deviation. (**F**) Grey inset: nascent leading strand products released by nicking with Nb.BsmI. pDPC^2xLead^ was replicated in non-depleted extract containing [α–^32^P]dATP and supplemented with DMSO or p97i, and buffer or 15 μM Ub-VS. Samples were processed and analyzed as in Figure 1C. (**G**) DNA samples as in Figure S1F (but lacking [α–^32^P]dATP) were withdrawn, and plasmid-associated proteins were recovered under non-stringent conditions as described in Figure 1D. (**H**) Model of CMG dynamics on pmeDPC^2xLead^. After CMGs bypass the two-leading strand DPCs, CMGs converge, as seen during replication termination, whereupon they are ubiquitylated (perhaps by CRL2^Lrr1^), and unloaded by p97. (**I**) pDPC^ICL^ reactions from Figure 1F (but lacking [α–^32^P]dATP) were subjected to stringent plasmid pulldown as in Figure S1D and analyzed for M.HpaII levels. (**J**) pICL or pDPC^ICL^ was replicated egg extract supplemented with p97i to prevent CMG unloading at the ICL and Ub-VS to prevent DPC proteolysis. After 110 minutes (red), when leading strands have approached the DPC, 250 μM free ubiquitin was added to the reaction to promote DPC proteolysis and enhance translesion synthesis past the DPC so that the leading strand can be extended to any CMG stalled at the ICL. DNA was purified and nicked with Nb.BsmI and separated on a denaturing polyacrylamide gel as in Figure 1C. (**K**) pDPC^Lead^ was replicated in undepleted extract containing [α–^32^P]dATP and Ub-VS, supplemented with buffer or Cdc7i (PHA767491) at zero or eight minutes after NPE extract addition, which initiates replication, and analyzed as in Figure 1C.

**Figure S2.**
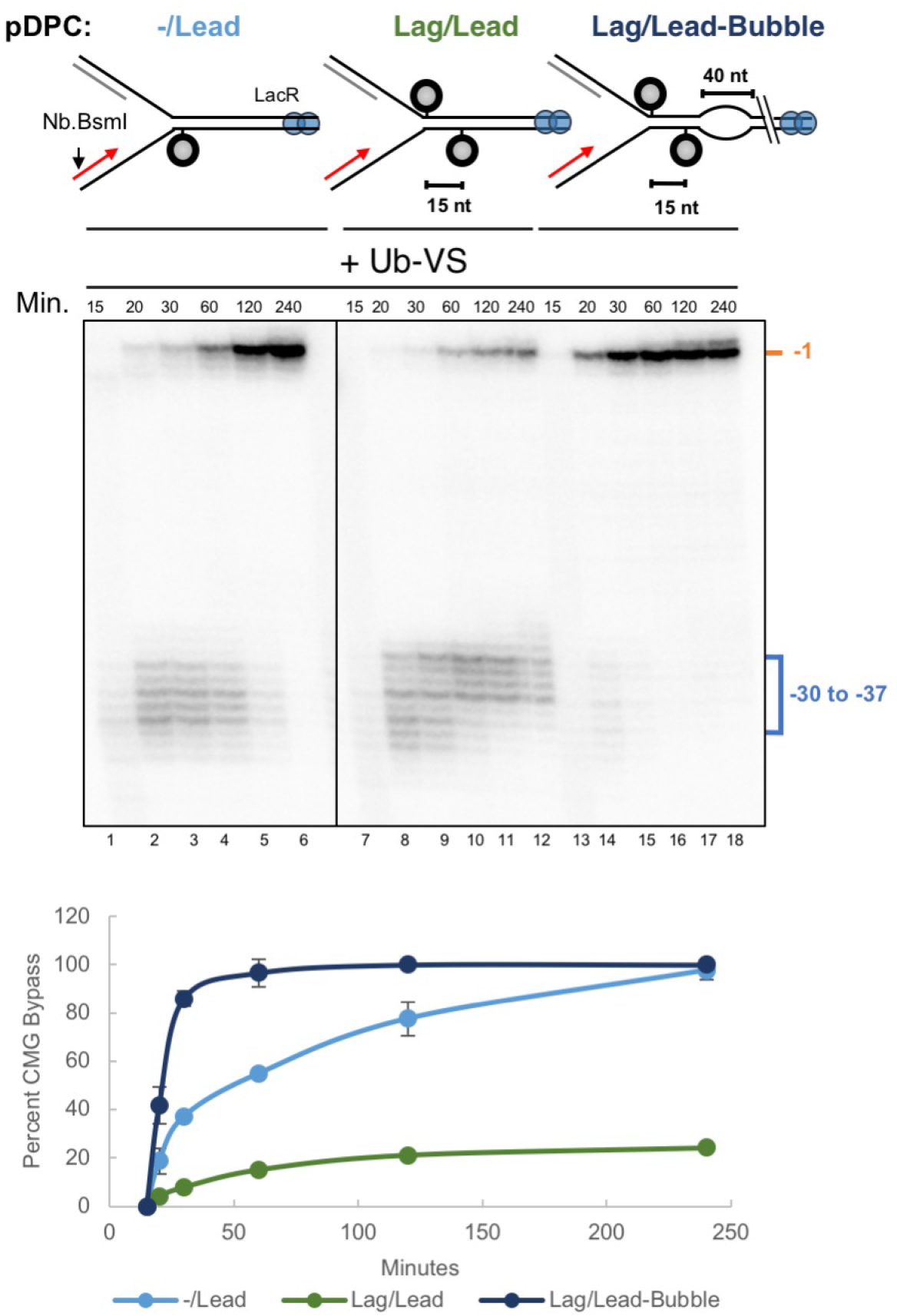
Related to Figure 2. pDPC^−/Lead^, pDPC^Lag/Lead^, and pmeDPC^Lag/Lead-Bubble^ were replicated in egg extracts containing [α–^32^P]dATP and 15 μM Ub-VS and analyzed as in Figure 1C. Approach of the nascent leading strands was used as a proxy for CMG bypass and quantified as in Figure 1C. The mean of three experiments is shown. Error bars represent the standard deviation.

**Figure S3.**
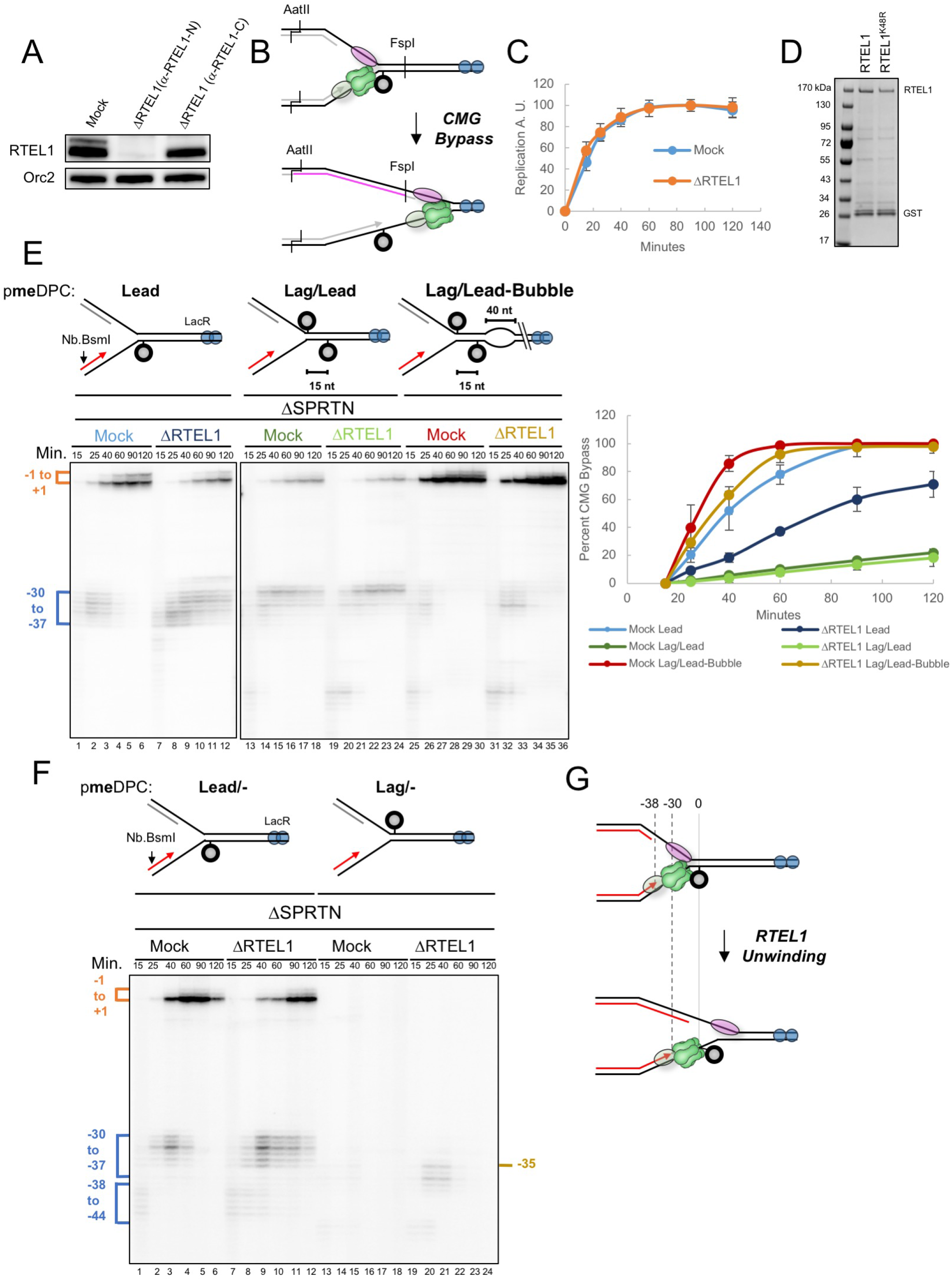
Related to Figure 3. (**A**) Egg extracts contain multiple RTEL1 isoforms, as seen in lane 1 and Figure 3B, lane 1. To distinguish them, we depleted egg extract with an antibody raised against an N-terminal fragment of RTEL1 (α-RTEL1-N, lane 2; used in all experiments except lane 3 of this panel) or an antibody raised against a C-terminal peptide (α-RTEL1-C, lane 3), and blotted with α-RTEL1-N or ORC2 (loading control) antibodies. Unlike α-RTEL1-N, α-RTEL1-C depleted only the largest RTEL1 isoform, consistent with the presence of multiple isoforms (as seen in mice (Ding et al., 2004)), only the largest of which has the C-terminal extension against which the antibody was raised. Depletion of extracts with α-RTEL1-C antibody (lane 3) had no effect on CMG bypass (data not shown), demonstrating that the shorter isoforms were sufficient to perform this function. (**B**) Schematic of lagging strand progression past the leading strand DPC following CMG bypass. The pink strand represents the lagging strand extension product generated after AatII and FspI digestion (shown in Figure 3D upper autoradiogram). (**C**) Incorporation of [α–^32^P]dATP during pmeDPC^Lead^ replication in mock-depleted or RTEL1-depleted extract was quantified and graphed. The mean of five experiments is shown. Error bars represent the standard deviation. (**D**) Coomassie blue-stained SDS-PAGE of purified RTEL1 wild-type and RTEL1-K48R. The RTEL1 and co-purifying GST-tag bands are indicated. (**E**) pmeDPC^Lead^, pmeDPC^Lag/Lead^, and pmeDPC^Lag/Lead-Bubble^ were replicated in the presence of [α–^32^P]dATP in SPRTN-depleted egg extracts that were also mock-depleted or RTEL1-depleted and analyzed as in Figure S1F. The mean of three experiments is quantified. Error bars represent the standard deviation. (**F**) Independent repeat of the experiment shown in Figure 3D, except that DNA samples were nicked with Nb.BsmI. As a result, no extension products were visible, and the percentage CMG bypass could not be quantified. (**G**) Model to account for the −38 arrest and how this is affected by RTEL1. When CMG first collides with a DPC, the duplex DNA underlying the DPC prevents CMG from reaching the adducted nucleotide, resulting in a −38 arrest. After unwinding of the DPC-associated DNA by RTEL1, CMG advances to the adducted nucleotide, yielding the −30 arrest.

**Figure S4.**
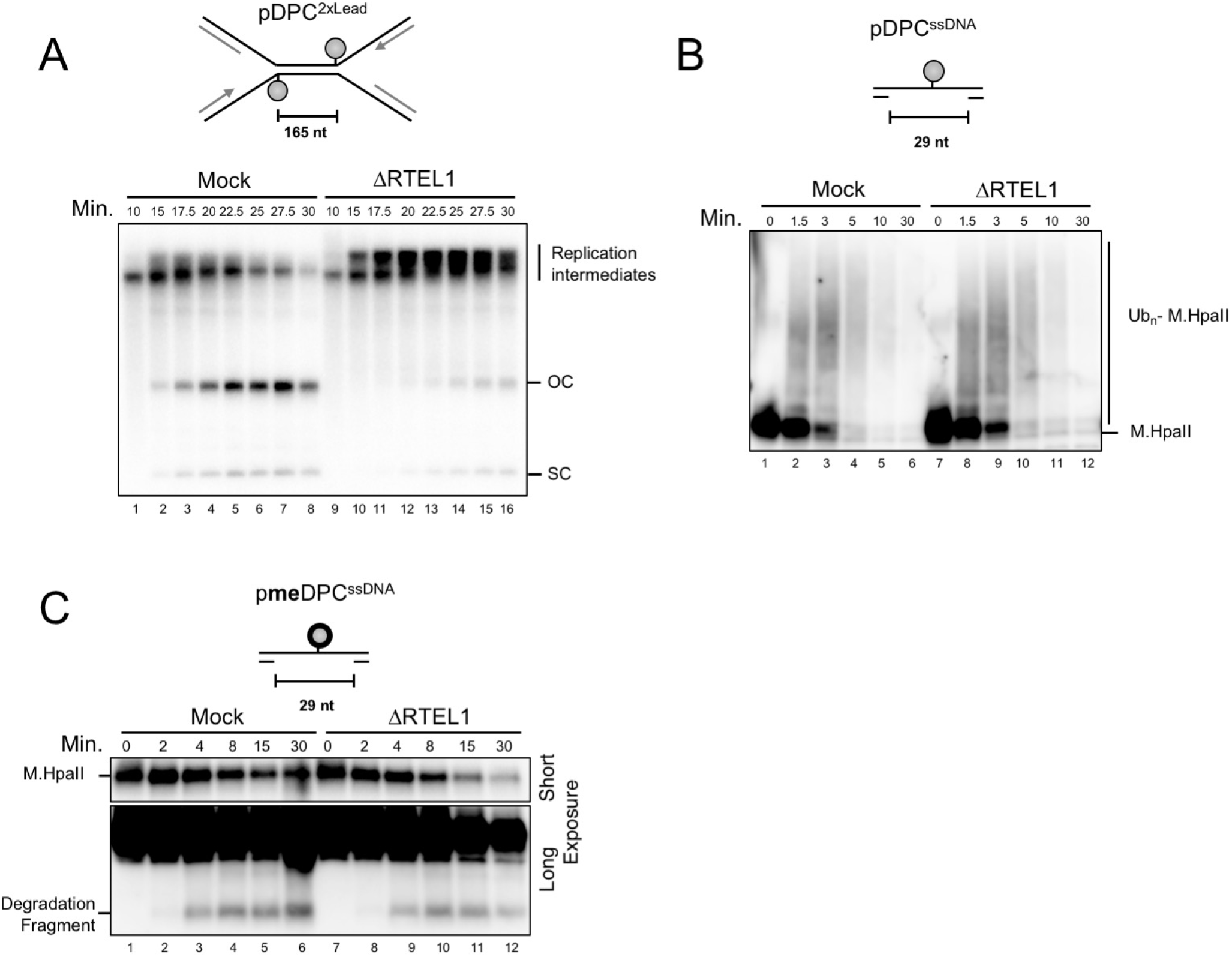
Related to Figure 5. (**A**) Mock-depleted and RTEL1-depleted extracts used in Figure 5C, Figure S4B, and Figure S4C were supplemented with [α–^32^P]dATP and used to replicate pDPC^2xLead^. DNA samples were separated on a native agarose gel and subjected to autoradiography. The gel shows that in the absence of RTEL1, replication products accumulated as slow mobility intermediates instead of undergoing dissolution into monomeric open circular (OC) and supercoiled (SC) plasmids, reflecting a defect in CMG bypass and lack of DNA unwinding of the 165 bp that separates the DPCs (see cartoon). (**B**) pDPC^ssDNA^ was incubated directly in mock-depleted or RTEL1-depleted NPE without first licensing in HSS, which prevents replication initiation due to the high concentration of Geminin in NPE (Arias and Walter, 2005). Chromatin was isolated by the stringent pull-down procedure described in Figure 4A and analyzed for M.HpaII levels. RTEL1 depletion had no effect on the rate of M.HpaII destruction. **(C)** Same as in (B) except that M.HpaII conjugated to the gapped plasmid was methylated to inhibit the proteasome pathway. Short and long (larger excerpt) exposures of the same blot are shown. The result shows that in the setting of ssDNA, the SPRTN-specific product was generated at the same rate whether or not RTEL1 was present

**Figure S5.**
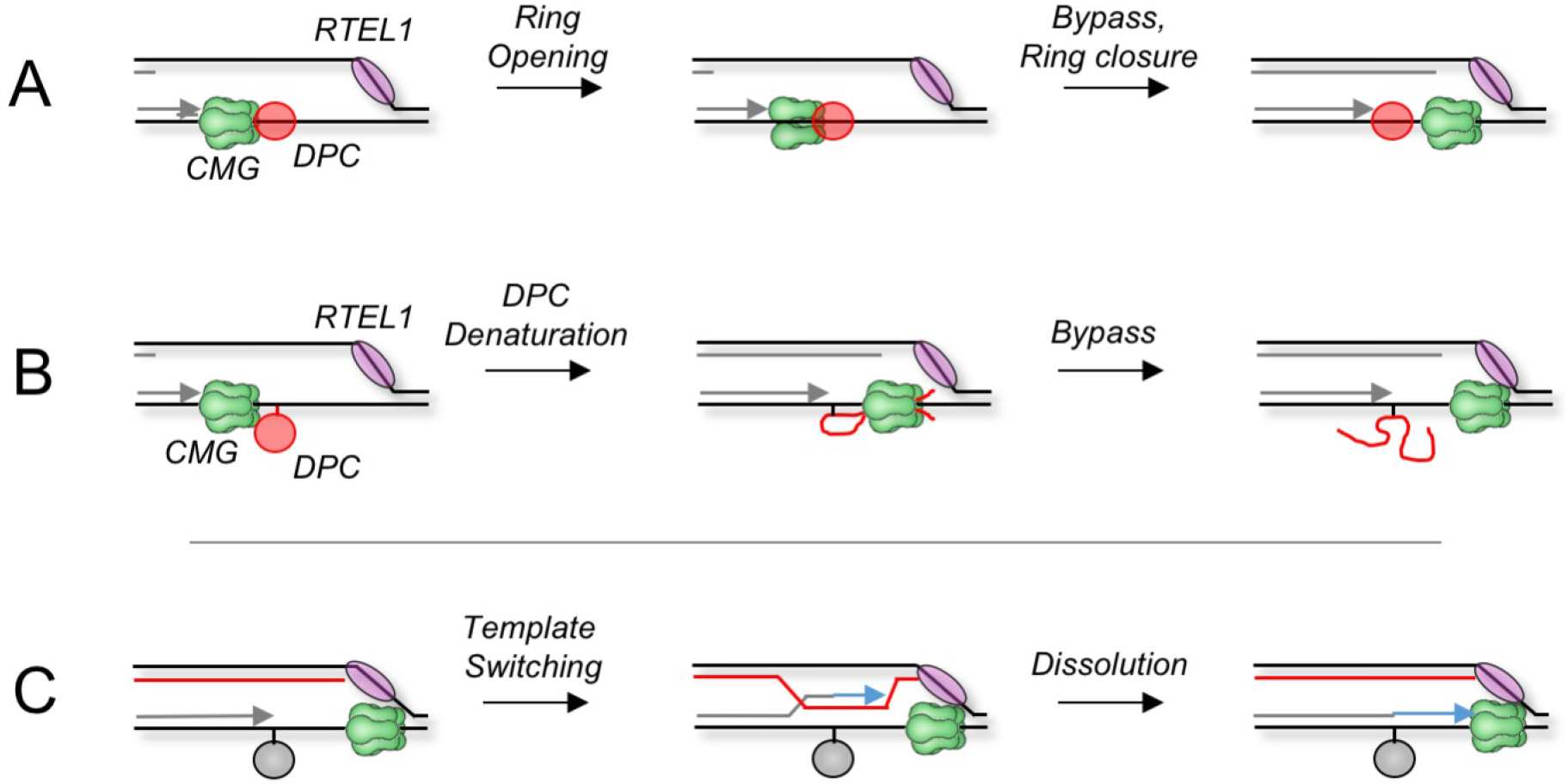
Related to Figure 7. Possible mechanisms of DPC bypass by CMG. **(A)** The MCM2-7 ring transiently opens, possibly due to dissociation of GINS and/or CDC45 (not shown), allowing the helicase to motor or be towed (not shown) past the DPC. **(B)** CMG denatures the DPC and threads the resulting polypeptide chains through the central channel of the MCM2-7. **(C)** Model for template switching at a DPC. If CMG bypasses the DPC but TLS fails, the leading strand anneals to the nascent lagging strand. After extension of the leading strand (blue arrow), it undergoes dissolution and re-anneals to the leading strand template, thereby bypassing the adduct.

## References

Arias, E.E., and Walter, J.C. (2005). Replication-dependent destruction of Cdt1 limits DNA replication to a single round per cell cycle in Xenopus egg extracts. Genes Dev 19, 114–126.

Bochman, M.L., and Schwacha, A. (2010). The Saccharomyces cerevisiae Mcm6/2 and Mcm5/3 ATPase active sites contribute to the function of the putative Mcm2-7 ‘gate’. Nucleic Acids Res 38, 6078–6088.

Brosh, R.M., Jr. (2013). DNA helicases involved in DNA repair and their roles in cancer. Nat Rev Cancer 13, 542–558.

Byun, T.S., Pacek, M., Yee, M.C., Walter, J.C., and Cimprich, K.A. (2005). Functional uncoupling of MCM helicase and DNA polymerase activities activates the ATR-dependent checkpoint. Genes Dev 19, 1040–1052.

Cali, F., Bharti, S.K., Di Perna, R., Brosh, R.M., Jr., and Pisani, F.M. (2016). Tim/Timeless, a member of the replication fork protection complex, operates with the Warsaw breakage syndrome DNA helicase DDX11 in the same fork recovery pathway. Nucleic Acids Res 44, 705–717.

Cantor, S.B., and Nayak, S. (2016). FANCJ at the FORK. Mutat Res 788, 7–11.

Chen, L., MacMillan, A.M., Chang, W., Ezaz-Nikpay, K., Lane, W.S., and Verdine, G.L. (1991). Direct identification of the active-site nucleophile in a DNA (cytosine-5)-methyltransferase. Biochemistry 30, 11018–11025.

Dewar, J.M., Budzowska, M., and Walter, J.C. (2015). The mechanism of DNA replication termination in vertebrates. Nature 525, 345–350.

Dewar, J.M., Low, E., Mann, M., Raschle, M., and Walter, J.C. (2017). CRL2(Lrr1) promotes unloading of the vertebrate replisome from chromatin during replication termination. Genes Dev 31, 275–290.

Ding, H., Schertzer, M., Wu, X., Gertsenstein, M., Selig, S., Kammori, M., Pourvali, R., Poon, S., Vulto, I., Chavez, E., et al. (2004). Regulation of murine telomere length by Rtel: an essential gene encoding a helicase-like protein. Cell 117, 873–886.

Douglas, M.E., Ali, F.A., Costa, A., and Diffley, J.F.X. (2018). The mechanism of eukaryotic CMG helicase activation. Nature.

Duxin, J.P., Dewar, J.M., Yardimci, H., and Walter, J.C. (2014). Repair of a DNA-protein crosslink by replication-coupled proteolysis. Cell 159, 346–357.

Enoiu, M., Ho, T.V., Long, D.T., Walter, J.C., and Scharer, O.D. (2012). Construction of plasmids containing site-specific DNA interstrand cross-links for biochemical and cell biological studies. Methods Mol Biol 920, 203–219.

Fu, Y.V., Yardimci, H., Long, D.T., Guainazzi, A., Bermudez, V.P., Hurwitz, J., van Oijen, A., Scharer, O.D., and Walter, J.C. (2011). Selective Bypass of a Lagging Strand Roadblock by the Eukaryotic Replicative DNA Helicase. Cell 146, 931–941.

Fullbright, G., Rycenga, H.B., Gruber, J.D., and Long, D.T. (2016). p97 Promotes a Conserved Mechanism of Helicase Unloading during DNA Cross-Link Repair. Mol Cell Biol 36, 2983–2994.

Gagou, M.E., Ganesh, A., Phear, G., Robinson, D., Petermann, E., Cox, A., and Meuth, M. (2014). Human PIF1 helicase supports DNA replication and cell growth under oncogenic-stress. Oncotarget 5, 11381–11398.

Garbelli, A., Beermann, S., Di Cicco, G., Dietrich, U., and Maga, G. (2011). A motif unique to the human DEAD-box protein DDX3 is important for nucleic acid binding, ATP hydrolysis, RNA/DNA unwinding and HIV-1 replication. PLoS One 6, e19810.

Georgescu, R., Yuan, Z., Bai, L., de Luna Almeida Santos, R., Sun, J., Zhang, D., Yurieva, O., Li, H., and O’Donnell, M.E. (2017). Structure of eukaryotic CMG helicase at a replication fork and implications to replisome architecture and origin initiation. Proc Natl Acad Sci U S A 114, E697–E706.

Guy, C.P., Atkinson, J., Gupta, M.K., Mahdi, A.A., Gwynn, E.J., Rudolph, C.J., Moon, P.B., van Knippenberg, I.C., Cadman, C.J., Dillingham, M.S., et al. (2009). Rep provides a second motor at the replisome to promote duplication of protein-bound DNA. Mol Cell 36, 654–666.

Hashimoto, Y., Puddu, F., and Costanzo, V. (2011). RAD51- and MRE11-dependent reassembly of uncoupled CMG helicase complex at collapsed replication forks. Nat Struct Mol Biol 19, 17–24.

Huang, J., Liu, S., Bellani, M.A., Thazhathveetil, A.K., Ling, C., de Winter, J.P., Wang, Y., Wang, W., and Seidman, M.M. (2013). The DNA Translocase FANCM/MHF Promotes Replication Traverse of DNA Interstrand Crosslinks. Molecular cell.

Ide, H., Shoulkamy, M.I., Nakano, T., Miyamoto-Matsubara, M., and Salem, A.M. (2011). Repair and biochemical effects of DNA-protein crosslinks. Mutat Res 711, 113–122.

Ivessa, A.S., Zhou, J.Q., and Zakian, V.A. (2000). The Saccharomyces Pif1p DNA helicase and the highly related Rrm3p have opposite effects on replication fork progression in ribosomal DNA. Cell 100, 479–489.

Klimasauskas, S., Kumar, S., Roberts, R.J., and Cheng, X. (1994). HhaI methyltransferase flips its target base out of the DNA helix. Cell 76, 357–369.

Langston, L., and O’Donnell, M. (2017). Action of CMG with strand-specific DNA blocks supports an internal unwinding mode for the eukaryotic replicative helicase. Elife 6.

Langston, L.D., Mayle, R., Schauer, G.D., Yurieva, O., Zhang, D., Yao, N.Y., Georgescu, R.E., and O’Donnell, M.E. (2017). Mcm10 promotes rapid isomerization of CMG-DNA for replisome bypass of lagging strand DNA blocks. Elife 6.

Lebofsky, R., Takahashi, T., and Walter, J.C. (2009). DNA replication in nucleus-free Xenopus egg extracts. Methods Mol Biol 521, 229–252.

Lessel, D., Vaz, B., Halder, S., Lockhart, P.J., Marinovic-Terzic, I., Lopez-Mosqueda, J., Philipp, M., Sim, J.C., Smith, K.R., Oehler, J., et al. (2014). Mutations in SPRTN cause early onset hepatocellular carcinoma, genomic instability and progeroid features. Nat Genet 46, 1239–1244.

Ling, C., Huang, J., Yan, Z., Li, Y., Ohzeki, M., Ishiai, M., Xu, D., Takata, M., Seidman, M., and Wang, W. (2016). Bloom syndrome complex promotes FANCM recruitment to stalled replication forks and facilitates both repair and traverse of DNA interstrand crosslinks. Cell Discov 2, 16047.

Lopez-Mosqueda, J., Maddi, K., Prgomet, S., Kalayil, S., Marinovic-Terzic, I., Terzic, J., and Dikic, I. (2016). SPRTN is a mammalian DNA-binding metalloprotease that resolves DNA-protein crosslinks. Elife 5.

Maric, M., Maculins, T., De Piccoli, G., and Labib, K. (2014). Cdc48 and a ubiquitin ligase drive disassembly of the CMG helicase at the end of DNA replication. Science 346, 1253596.

Maskey, R.S., Flatten, K.S., Sieben, C.J., Peterson, K.L., Baker, D.J., Nam, H.J., Kim, M.S., Smyrk, T.C., Kojima, Y., Machida, Y., et al. (2017). Spartan deficiency causes accumulation of Topoisomerase 1 cleavage complexes and tumorigenesis. Nucleic Acids Res 45, 4564–4576.

Maskey, R.S., Kim, M.S., Baker, D.J., Childs, B., Malureanu, L.A., Jeganathan, K.B., Machida, Y., van Deursen, J.M., and Machida, Y.J. (2014). Spartan deficiency causes genomic instability and progeroid phenotypes. Nat Commun 5, 5744.

Mimura, S., and Takisawa, H. (1998). Xenopus Cdc45-dependent loading of DNA polymerase alpha onto chromatin under the control of S-phase Cdk. Embo J 17, 5699–5707.

Moreno, S.P., Bailey, R., Campion, N., Herron, S., and Gambus, A. (2014). Polyubiquitylation drives replisome disassembly at the termination of DNA replication. Science 346, 477–481.

Morocz, M., Zsigmond, E., Toth, R., Enyedi, M.Z., Pinter, L., and Haracska, L. (2017). DNA-dependent protease activity of human Spartan facilitates replication of DNA-protein crosslink-containing DNA. Nucleic Acids Res 45, 3172–3188.

Nakano, T., Miyamoto-Matsubara, M., Shoulkamy, M.I., Salem, A.M., Pack, S.P., Ishimi, Y., and Ide, H. (2013). Translocation and stability of replicative DNA helicases upon encountering DNA-protein cross-links. J Biol Chem 288, 4649–4658.

O’Donnell, M.E., and Li, H. (2018). The ring-shaped hexameric helicases that function at DNA replication forks. Nat Struct Mol Biol 25, 122–130.

Raschle, M., Knipsheer, P., Enoiu, M., Angelov, T., Sun, J., Griffith, J.D., Ellenberger, T.E., Scharer, O.D., and Walter, J.C. (2008). Mechanism of replication-coupled DNA interstrand crosslink repair. Cell 134, 969–980.

Reinisch, K.M., Chen, L., Verdine, G.L., and Lipscomb, W.N. (1995). The crystal structure of HaeIII methyltransferase convalently complexed to DNA: an extrahelical cytosine and rearranged base pairing. Cell 82, 143–153.

Rosado, I.V., Langevin, F., Crossan, G.P., Takata, M., and Patel, K.J. (2011). Formaldehyde catabolism is essential in cells deficient for the Fanconi anemia DNA-repair pathway. Nature structural & molecular biology 18, 1432–1434.

Samel, S.A., Fernandez-Cid, A., Sun, J., Riera, A., Tognetti, S., Herrera, M.C., Li, H., and Speck, C. (2014). A unique DNA entry gate serves for regulated loading of the eukaryotic replicative helicase MCM2-7 onto DNA. Genes & development 28, 1653–1666.

Semlow, D.R., Zhang, J., Budzowska, M., Drohat, A.C., and Walter, J.C. (2016). Replication-Dependent Unhooking of DNA Interstrand Cross-Links by the NEIL3 Glycosylase. Cell 167, 498–511 e414.

Stingele, J., Bellelli, R., Alte, F., Hewitt, G., Sarek, G., Maslen, S.L., Tsutakawa, S.E., Borg, A., Kjaer, S., Tainer, J.A., et al. (2016). Mechanism and Regulation of DNA-Protein Crosslink Repair by the DNA-Dependent Metalloprotease SPRTN. Mol Cell 64, 688–703.

Stingele, J., Bellelli, R., and Boulton, S.J. (2017). Mechanisms of DNA-protein crosslink repair. Nat Rev Mol Cell Biol 18, 563–573.

Stingele, J., Schwarz, M.S., Bloemeke, N., Wolf, P.G., and Jentsch, S. (2014). A DNA-dependent protease involved in DNA-protein crosslink repair. Cell 158, 327–338.

Vannier, J.B., Sandhu, S., Petalcorin, M.I., Wu, X., Nabi, Z., Ding, H., and Boulton, S.J. (2013). RTEL1 is a replisome-associated helicase that promotes telomere and genome-wide replication. Science 342, 239–242.

Vannier, J.B., Sarek, G., and Boulton, S.J. (2014). RTEL1: functions of a disease-associated helicase. Trends Cell Biol 24, 416–425.

Vaz, B., Popovic, M., Newman, J.A., Fielden, J., Aitkenhead, H., Halder, S., Singh, A.N., Vendrell, I., Fischer, R., Torrecilla, I., et al. (2016). Metalloprotease SPRTN/DVC1 Orchestrates Replication-Coupled DNA-Protein Crosslink Repair. Mol Cell 64, 704–719.

Vaz, B., Popovic, M., and Ramadan, K. (2017). DNA-Protein Crosslink Proteolysis Repair. Trends Biochem Sci 42, 483–495.

Walter, J., and Newport, J.W. (1997). Regulation of replicon size in Xenopus egg extracts. Science 275, 993–995.

Yardimci, H., Wang, X., Loveland, A.B., Tappin, I., Rudner, D.Z., Hurwitz, J., van Oijen, A.M., and Walter, J.C. (2012). Bypass of a protein barrier by a replicative DNA helicase. Nature 492, 205–209.

Yuan, Z., Bai, L., Sun, J., Georgescu, R., Liu, J., O’Donnell, M.E., and Li, H. (2016). Structure of the eukaryotic replicative CMG helicase suggests a pumpjack motion for translocation. Nat Struct Mol Biol 23, 217–224.

